# Mercury methylation trait dispersed across diverse anaerobic microbial guilds in a eutrophic sulfate-enriched lake

**DOI:** 10.1101/2020.04.01.018762

**Authors:** Benjamin D. Peterson, Elizabeth A. McDaniel, Anna G. Schmidt, Ryan F. Lepak, Patricia Q. Tran, Robert A. Marick, Jacob M. Ogorek, John F. DeWild, David P. Krabbenhoft, Katherine D. McMahon

## Abstract

Mercury (Hg) methylation is a microbially mediated process that converts inorganic Hg into the bioaccumulative neurotoxin methylmercury (MeHg). Exploring the diversity and metabolic potential of the dominant Hg-methylating microorganisms can provide insights into how biogeochemical cycles and water quality parameters underlie MeHg production. However, our understanding of the ecophysiology of methylators in natural ecosystems is still limited. Here, we used shotgun metagenomics paired with biogeochemical data to identify likely hotspots for MeHg production in a lake with elevated sulfate levels and characterize the microbial methylators and the flanking microbial community. Identified putative methylators were dominated by hgcA sequences divergent from those in canonical, experimentally confirmed methylators. Using genome-resolved metagenomics, these sequences were identified within genomes associated with Bacteroidetes and the recently described phylum Kiritimatiellaeota. Over half of the hgcA abundance comes from genomes corresponding to obligately fermentative organisms, many of which have a large number of glucoside hydrolases for polysaccharide degradation. Sulfate-reducing genomes encoding hgcA were also identified, but only accounted for 22% of the abundance of hgcA+ genomes. This work highlights the diverse dispersal of the methylation trait across the microbial anoxic food web.

## Introduction

Mercury (Hg) contamination of aquatic food webs is an environmental concern and a public health hazard. Environmental levels of Hg have increased drastically due to anthropogenic inputs, such as burning coal for electricity and artisanal gold mining.^1^ Much of this anthropogenic Hg is in the form of elemental (Hg(0)) or inorganic (Hg(II)) Hg.^2^ However, Hg bioaccumulates in tissues and biomagnifies up food webs in the form of methylmercury (MeHg), making the production of MeHg the gateway process to food web contamination.^3^ MeHg production is mediated by microorganisms in aquatic anoxic environments such as sediments, periphyton, rice paddy soils, and the freshwater and marine water column.^4–8^ MeHg accumulation in freshwater hypolimnia has historically been attributed to production in the sediment and diffusion across the sediment-water interface.^5, 9,10^ However, methylation has been shown to occur in the water column and may account for a substantial fraction of the MeHg hypolimnetic accumulation, especially in lakes with a large anoxic hypolimnion.^8, 11–13^ Despite this, water column methylation in freshwater lakes is understudied relative to sediment methylation.

The production of MeHg is driven largely by local geochemical conditions. Sulfide levels and the quality and quantity of dissolved organic matter (DOM) impact the complexation and aggregation of Hg(II).^14, 15^ This has a direct impact on MeHg production by controlling the bioavailability of Hg to methylating organisms. Additionally, the identity and quantity of electron acceptors and donors can indirectly drive MeHg production by fueling the metabolic activity of methylating microorganisms.^10, 16^ The identification and characterization of these organisms can illuminate how biogeochemical processes are influencing MeHg production. Early experiments on cultured isolates and *in situ* assays showing molybdate inhibition of MeHg production in sediments provided a link between sulfate-reducing bacteria (SRBs) and the production of MeHg.^9, 10^ Later studies identified several iron-reducing bacteria (FeRBs) and methanogenic archaea as methylators as well.^17, 18^ More generally, MeHg production often increases with increasing overall heterotrophic activity, suggesting that increased carbon and energy flux through the microbial food web can drive MeHg production.^8, 19^

The recent identification of the *hgcAB* gene cluster has provided a robust molecular marker for methylation potential.^20^ This marker has been used to search publicly available genomes, metagenomes and metagenome-assembled genomes, which has expanded the phylogenetic and metabolic diversity of confirmed and putative methylators.^21–25^ Several PCR primer sets have been developed to identify *hgcA* sequences in natural systems.^26–30^ Studies using this approach in habitats such as rice paddies, soils/sediments, periphyton, and many others have shown that the hgcA+ community can be quite different phylogenetically across environments, and that this can sometimes be linked to biogeochemical conditions at the site.^26, 29–32^ While this approach works well in some systems, these primers do not always accurately characterize the hgcA+ community in natural ecosystems, especially for *hgcA* sequences that are highly divergent compared to reference methylators.^12, 23, 33, 34^ Shotgun metagenomics offers a more robust method of identifying *hgcA* sequences from environmental samples, since computational tools such as BLAST and Hidden Markov Models (HMMs) are better equipped to identify divergent sequences.^35^ For example, Gionfriddo et al used an assembly-based metagenomic approach to identify a divergent *hgcA* sequence fragment from the Antarctic sea ice and determined it was likely from a nitrite-oxidizing *Nitrospina*.^36^ While this approach improves gene detection, it does not provide key information on the metabolic capabilities of hgcA+ organisms and relies on relatively small databases with relevant sequences for comparison. As an alternative, genome-resolved metagenomics yields population genomes (bins) from complex environmental samples.^37^ This approach enables phylogenetic identification of hgcA+ lineages and characterization of their metabolic potential. Recently, Jones et al. applied this approach in two sulfate-enriched lakes and identified a broad diversity of hgcA+ groups, including some previously unknown methylators, and metabolically characterized each of these draft genomes.^12^

The primary objective of this study was to identify and characterize the Hg-methylating microorganisms in the anoxic hypolimnion of a eutrophic, sulfate-enriched lake. We monitored the biogeochemical and redox status of the hypolimnion throughout the ice-free season and generated Hg speciation profiles. We selected samples for shotgun metagenomic sequencing from sites where we suspected *in situ* MeHg production. We used assembly-based and bin-based analyses to characterize the phylogenetic diversity and metabolic potential of the hgcA+ microbial community. We show that the hgcAB+ community in Lake Mendota is dominated by non-canonical methylators and that fermentative organisms account for over half of the hgcAB+ population, but that microorganisms throughout the anaerobic food web are potentially involved, either directly or indirectly, in MeHg production.

## Materials and Methods

Detailed methods can be found in the Supplementary Methods online.

### Field sampling

Lake Mendota is a large, dimictic lake located in Madison, Wisconsin, USA. Sampling was conducted within 100m of the North Temperate Lakes Long-Term Ecological Research buoy (GPS coordinates: 43.09885, −89.40545) at the Deep Hole, the deepest basin in Lake Mendota. Samples were collected approximately monthly in 2017 from the onset of anoxia until turnover. Detailed profiles of temperature, dissolved oxygen and turbidity were collected with a YSI Exo2 multiparameter sonde (YSI Incorporated, Yellow Springs, OH). These profiles were viewed in real-time to guide sampling. All samples were collected through an acid-washed Teflon sampling line using a peristaltic pump. Samples for sulfide analysis were preserved in 1% ZnAc. Water samples for dissolved metal analysis were filtered through a 0.45µm PES Acrodisc filter and acidified to 1% HCl. Hg samples were collected using clean hands/dirty hands. Samples were collected into a new PETG 2.5L bottle, which was allowed to overflow before capping. Hg samples were double-bagged and stored in a cooler, then at 4°C in the dark. Water was filtered through a quartz fiber filter (QFF) within 24 hours and preserved with 1% HCl for dissolved Hg species analysis. The filters were frozen for particulate Hg analysis. DNA samples were collected onto 0.22µm pore-size PES filters (Pall Corp.) and flash-frozen on liquid nitrogen within 90 seconds.

### Geochemical analyses

Sulfide was quantified spectrophotometrically using the Cline method.^38^ Iron and manganese were quantified by inductively-coupled plasma optical emission spectrometry (ICP-OES) on a Varian Vista-MPX CCD ICP-OES. Processing and analysis of Hg samples was done at the U.S. Geological Survey (USGS) Wisconsin Mercury Research Laboratory. Dissolved total Hg (THg) was quantified after purge and trap using cold vapor atomic fluorescence spectrometry, using a Tekran Model 2500 CVAFS Mercury Detector (Tekran Instruments Corps., Toronto, ON, Canada). Particulate THg was extracted from the filter using 5% bromium chloride before purge and trap. This protocol follows the U.S. Environmental Protection Agency (EPA) Method 1631. Dissolved MeHg was quantified using isotope dilution by distillation, gas chromatography separation and inductively-coupled plasma-mass spectrometry (ICP-MS) on a Thermo ICAP-RQ ICP-MS (Thermo).

### DNA extraction, sequencing, and assembly

DNA was extracted by enzymatic and physical lysis followed by phenol-chloroform extraction and purification by isopropanol precipitation. DNA library preparation was done at the Functional Genomics Lab and sequencing was done in the Vincent J. Coates Genomics Sequencing Lab, both within the California Institute for Quantitative Biosciences (QB3-Berkeley, Berkeley, CA). Library preparation was done with a Kapa Biosystem Library Prep kit, targeting inserts ∼600bp in length (Roche Sequencing and Life Science, Kapa Biosystems, Wilmington, MA). Libraries were pooled and sequenced on a single lane of an Illumina HiSeq4000 for paired-end reads of 150bp (Illumina, Inc., San Diego, CA). Raw reads were trimmed using Sickle to maintain a QC score of 20 over a sliding window of 15.^39^ Trimmed reads shorter than 100bp were cut. Metagenomes were both assembled individually and co-assembled using metaSPADes (v3.12).^40^ Assembly-based analyses were run on all scaffolds at least 500bp in length.

### Metagenomic analysis, binning and annotation

A custom HMM for HgcA amino acid sequences was built using hmmbuild from hmmer (v3.1b2) using experimentally verified HgcA amino acid sequences (Data File 2).^21, 41^ This HMM was used to identify HgcA sequences from the open reading frames of each assembly using the trusted cutoff score of 128.60. Each putative HgcA sequence was manually screened and discarded if it did not contain the cap helix domain (N(V/I)WCA(A/G)(A/G)(K/R)) and at least 4 transmembrane domains (Figure S2).^20^ We dereplicated the HgcA sequences across assemblies using CD-HIT, clustering them at 0.97 identity.^42^ Reads from all metagenomes were mapped to the scaffolds from each assembly individually using BBMap (v35) with default settings.^43^ Open reading frames were predicted using Prodigal (v2.6.2).^44^ Automatic binning was done individually on each assembly, using only scaffolds >1000bp in length. Bins were generated using Metabat2, MaxBin (v2.1.1), and CONCOCT (v0.4.1), then aggregated using Das Tool.^45–48^ Bins across assemblies were clustered into “high matching sets” (HMSs) if they shared at least 98% ANI over at least 50% of the genome. CheckM was used to estimate the completeness and redundancy of each bin.^49^ One bin from each HMS was selected for analysis, based primarily on percent completeness, and quality of the assembly. Bins were then decontaminated using the anvi-refine interface in Anvi’o (v5.2).^50^ All hgcA+ bins were reassembled using SPADes and manually re-binned in Anvi’o. Manual comparison of the GC content, tetranucleotide frequency, differential coverage, and taxonomy of adjacent genes was conducted on binned hgcA+ scaffolds, relative to other scaffolds in the bin, to confirm the inclusion of these scaffolds within the bin. Taxonomy of each bin was automatically assigned using the GTDB-TK software.^51^ Preliminary annotations were done using MetaPathways.^52^ Annotations of metabolic genes of interest were confirmed using Hidden Markov Models (HMMs) from TIGRFAM and PFAM.^35^ In many cases, gene neighborhoods and phylogenies were also used to confirm annotations.

### Phylogenetic analyses

Bin phylogenies were based on 16 ribosomal protein sequences.^53^ For both bin and HgcA phylogenies, amino acid sequences were aligned using MUSCLE (v3.8.31).^54^ For the bin alignment, all 16 rp16 gene alignments were concatenated into a single alignment. Sequences with less than half of the aligned residues were manually removed. Alignments were manually inspected in Geneious and trimmed using BMGE1.1 with the BLOSUM30 substitution matrix.^55^ RAxML (v8.2.11) was used to generate a maximum likelihood (ML) tree under the GAMMA distribution with the LG model.^56^ Branch support was generated by rapid bootstrapping. For HgcA phylogenies, we used RogueNaRok (v1.0) to identify and remove “rogue taxa” interfering with proper tree generation.^57^ The best-scoring ML tree for HgcA was mid-point rooted using the Phangorn R package and visualized using ggtree.^58, 59^ The rp16 ML tree was rooted using an archaeal outgroup and visualized using ggtree.

## Results and Discussion

### Hg and redox biogeochemistry in Lake Mendota

Lake Mendota is a eutrophic lake enriched in sulfate, with a watershed dominated by agriculture, leading to high levels of nutrient inflow and productivity. Physical and biogeochemical profiles were collected approximately monthly over the stratified period in 2017. A subset of the profiles are shown in Figure 1 and S1. Anoxia developed in the hypolimnion as early as June, likely due to the intense spring blooms sinking and decomposing. Reduced iron (Fe) and manganese (Mn) accumulated in the hypolimnion immediately following anoxia onset. While the Fe was quickly precipitated out by sulfide, Mn continued to accumulate in the hypolimnion up to ∼5µM.^60^ We observed an enrichment of dissolved and particulate Mn near the oxic/anoxic interface during late stratification (September and October), suggesting enhanced redox cycling in this region (particulate data not shown). Mendota has relatively high sulfate concentrations, with up to ∼175-200µM in the epilimnion. Combined with early anoxia and continued primary production in the epilimnion, this provides a rich habitat for sulfate reduction, which has previously been shown to occur in both the sediments and the water column.^61^ Sulfide was detectable within 1 meter below the oxycline as early as August and accumulated to over 150µM by October.

**Figure 1.**
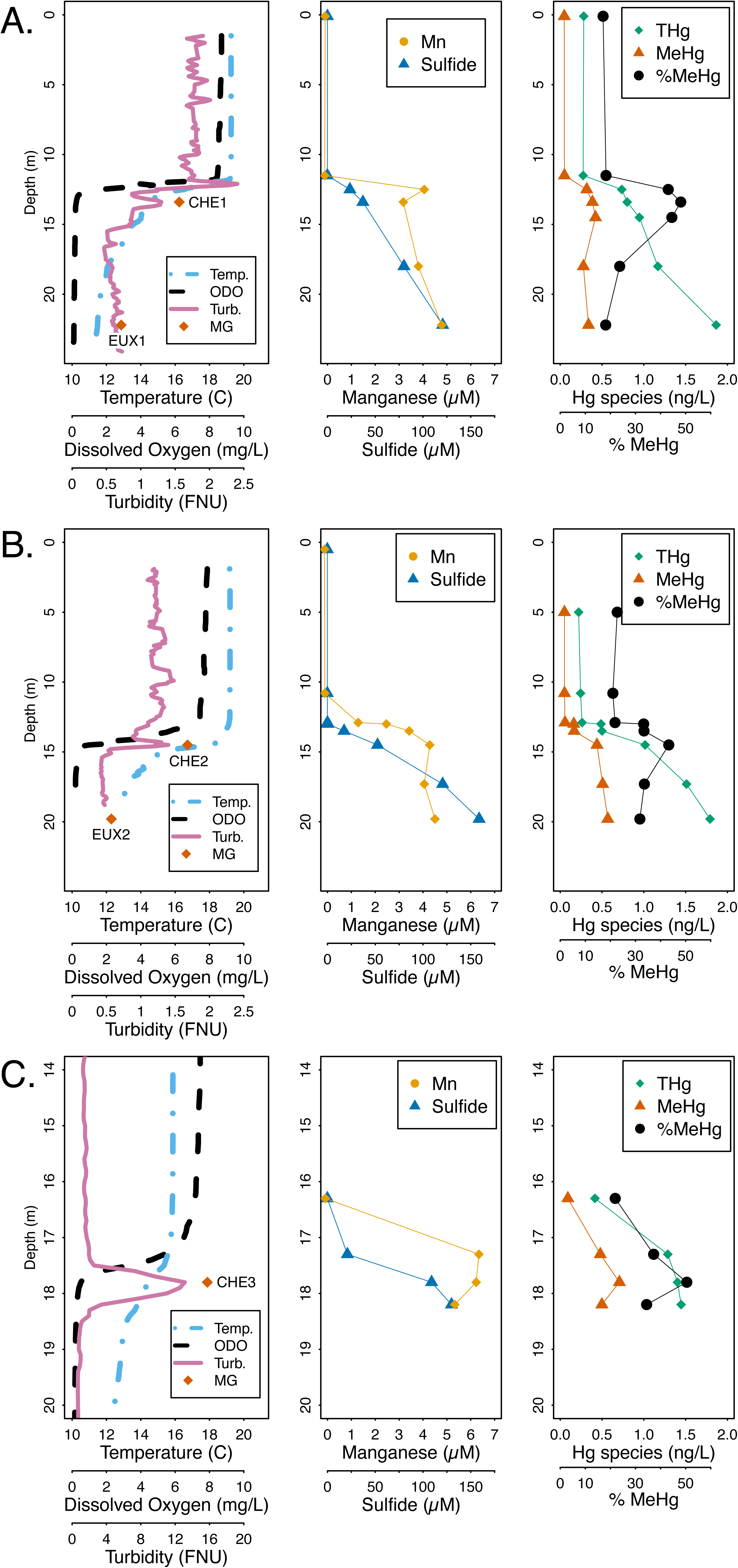
Physical and geochemical profiles of Lake Mendota from 2017 on September 8th (A), October 4th (B) and October 19th (C). The first column displays parameters measured continuously with an Exo2 sonde and includes orange diamonds where samples for metagenomic sequencing were collected. Names of the metagenomes are displayed near the orange diamonds. Equipment failure resulted in a slightly truncated sonde profile on October 4th (B). The second column displays sulfide and filter-passing manganese values at discrete depths. The total and methylmercury measurements, in the third column, are a bulk value, representing the sum total of the dissolved and particulate fractions. Dissolved and particulate fractions are plotted individually in Figure S1. Note the changed scale for depth on the y-axis and for turbidity on the x-axis in the October 19th profiles (C). The metagenomic samples collected near the metalimnion for October 4th and October 19th were both collected coincident with the observed spike in turbidity. Abbreviations: Temp. - Temperature (°C), ODO - Optical dissolved oxygen in mg/L, Turb. - Turbidity in Formazin Nephelometric Units (FNU), MG - metagenome sample, Mn - Filter-passing manganese, THg - Total mercury, MeHg - Methylmercury, %MeHg - Methylmercury concentration divided by total mercury concentration.

Once oxygen was depleted, both THg and (MeHg) began to accumulate in the hypolimnion (Figure 1, Figure S1). THg in the hypolimnion increased throughout the summer, starting at ∼0.5ng/L in the epilimnion and increasing down the water column. MeHg and THg continued to increase in the hypolimnion in August. In September and October, THg continued to increase in concentration, reaching nearly 2ng/L in the bottom waters. However, the MeHg increased across the oxic/anoxic interface, then stayed approximately even or decreased slightly with increasing depth. Correspondingly, the fraction MeHg relative to THg peaked at the oxic/anoxic interface. This mid-column peak of fraction MeHg is an indication of *in situ* production. This coincided with a peak in turbidity, which has been previously shown to co-localize with elevated microbial activity and MeHg production.^13^

### HgcA identification

To identify potential hgcA+ groups in Lake Mendota, we selected five samples for shotgun metagenomic DNA sequencing and analysis (Figure 1, Table S1). Three of these samples were selected to coincide with the mid-column peak in MeHg on three different dates. These chemocline samples will be referred to as CHE1, CHE2, and CHE3, based on their temporal order. The other two samples were collected from deeper, more euxinic (oxygen-depleted and sulfide-rich) waters on two separate dates. These euxinic samples will be referred to as EUX1 and EUX2, also based on their temporal order. Notably, the water from which CHE3 was sampled was relatively high in sulfide despite its proximity to the oxic-anoxic interface (Figure 1C). Metagenomes were assembled (statistics in Table S2) and binned (bin information in Data File 1). We retrieved 228 bins that were more than 75% complete and less than 10% redundant, which accounted for only 33% of the total number of reads.

We identified 108 unique HgcA sequences in the unbinned metagenomic assemblies using a custom-built HgcA HMM (Data File 2). The HgcA amino acid sequences are in Data File 3, and the nucleic acid files in Data File 4. Each identified HgcA was manually screened for the cap helix domain and at least 4 transmembrane domains (Figure S2)^20^. Ninety of the corresponding *hgcA* genes had a putative *hgcB* sequence downstream. Seven of the 18 hgcA+ scaffolds lacking *hgcB* ended just downstream of *hgcA*, and it is possible that *hgcB* simply did not assemble into the scaffold. The remaining 11 *hgcA* genes without an *hgcB* partner had a similar phylogenetic and coverage distribution to those with a downstream *hgcB* (Figure S3, Data File 5). Methylation has been experimentally verified in *Desulfovibrio africanus* sp. Walvis Bay, in which *hgcA* is separated from *hgcB* by a single ORF.^20, 62^ While there are no other studies on the methylation capacity of *hgcA* genes without a downstream *hgcB* that we know of, we included all identified sequences in our analysis for completeness.

We also searched for *hgcA* in the bins and discovered 41 hgcA+ bins. Manual comparisons of the GC content, tetranucleotide frequency, differential coverage, and adjacent gene phylogenies were conducted on binned hgcA+ scaffolds, relative to other scaffolds in the bin, to gather more evidence in support of the inclusion of these sequences within the bin. One of these bins (LEN_0031) included two copies of the *hgcA* gene. However, bins represent composite population genomes rather than individual genomes.^63^ Thus, we cannot confirm that the two *hgcA* sequences were present together in a single organism. These 41 bins accounted for 51% of the total *hgcA* coverage in our assemblies. This limited coverage highlights an inability to recover quality genomes harboring the most abundant *hgcA* genes. For example, 13 out of the 30 most abundant sequences were not binned even though they were on relatively long scaffolds (Figure S4). Efforts to recover highly abundant hgcA+ bins through read subsampling, contig curation using assembly graphs, reassembly, and manual binning and curation were unable to recover these highly abundant populations.

Despite the constraints of population genome recovery, our hgcA+ bins were representative of the overall *hgcA* diversity. We successfully binned contigs from most of the HgcA phylogenetic clusters identified using the unbinned contigs (Figure 2). The hgcA+ bins accounted for 17% of the total read coverage from all bins, and included some of the most abundant bins in our metagenomes (Figure S5a). They had slightly less coverage per bin than hgcA-bins (not significant), but this could be due to the greater degree of manual curation of the hgcA+ bins (Figure 5b). Overall, the hgcA+ bins recruited 6% of the total number of reads from our metagenomic datasets. Because the hgcA+ bins accounted for only 51% of the total coverage of all recovered *hgcA* sequences, we estimate that hgcA+ genomes account for ∼12% of the total metagenomic reads across our five samples.

**Figure 2.**
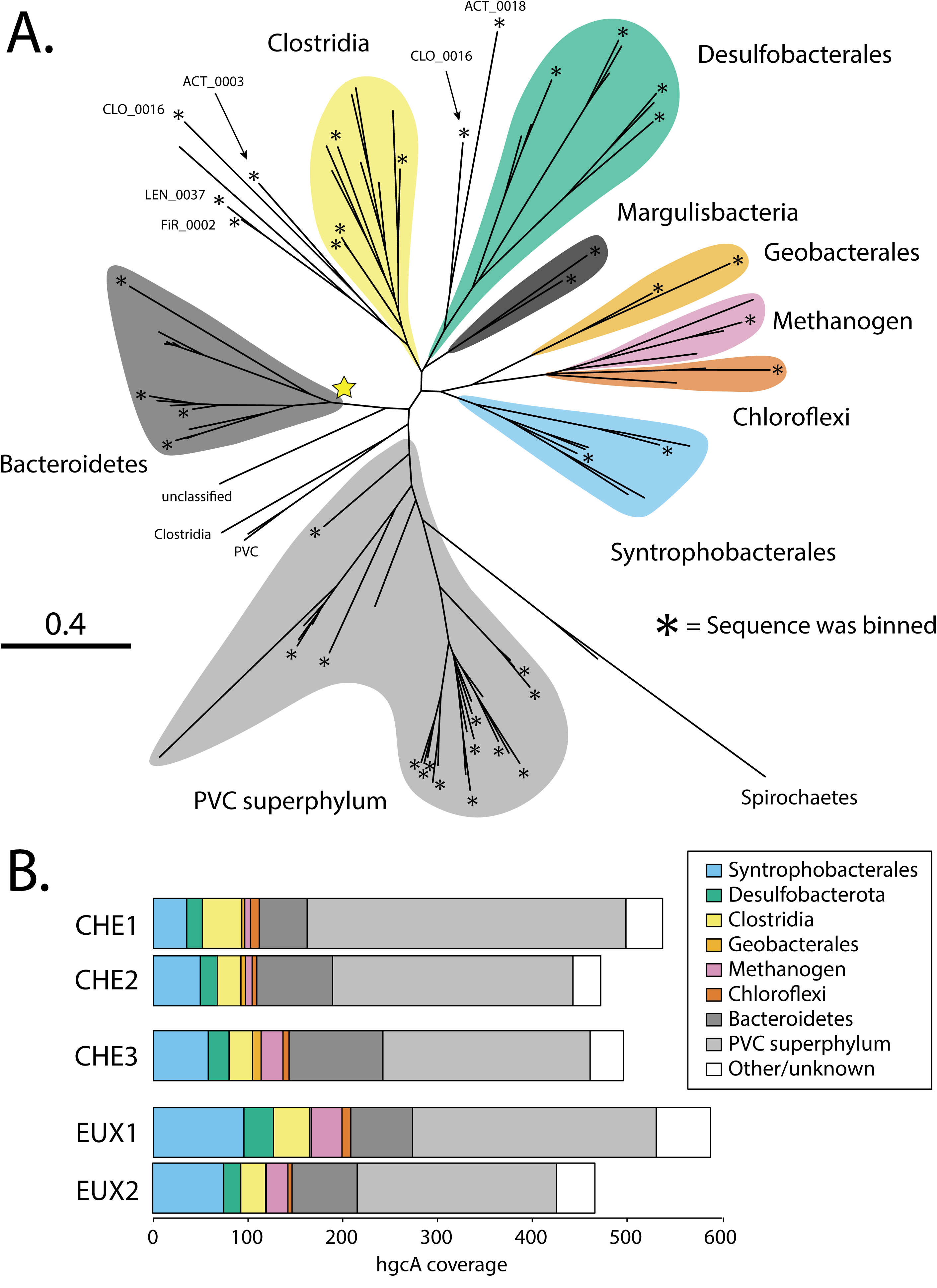
Unconfirmed methylators dominate hgcA sequence diversity in Lake Mendota, both numerically (A) and by coverage (B). A: Phylogenetic tree of 108 hgcA sequences from this study. Asterisks at the end of branches indicate sequence was binned. All other branches are unbinned hgcA sequences from this study. Sequences were assigned a predicted taxonomic group based on phylogenetic clustering with hgcA reference sequences from NCBI and bin phylogenies of binned hgcA sequences (for detailed tree with reference sequences, see Figure S3). Binned sequences outside of a monophylogenetic cluster are labeled with their bin name. The yellow star indicates the branch to the monophyletic Bacteroidetes sequences. B: Sum of coverage of hgcA sequences within predicted taxonomic groups across 5 metagenomic samples. Coverage refers to the average depth of coverage across hgcA+ scaffolds.

### Phylogenetic diversity of hgcA+ community

Of the 108 HgcA sequences, 43 of them clustered with experimentally verified HgcA sequences in the HgcA phylogenetic tree and accounted for 27% of the total coverage (Figure 2). Some of these sequences (21 sequences, 17% of total coverage) clustered with HgcA sequences from Deltaproteobacteria genomes. These included three major groups associated with three different microbial orders: Desulfobacterales (5% of total *hgcA* coverage), Geobacterales (<1%), and Syntrophobacterales (∼12%). No sequences associated with Desulfovibrionales, the order including the well-studied sulfate-reducing *Desulfovibrio desulfuricans* ND132, were detected.^64^ This relatively low number of *hgcA* genes associated with Deltaproteobacteria is not due to a predominance of hgcA-organisms from this class; rather, we only retrieved 15 total bins associated with Deltaproteobacteria, 9 of which were hgcA+. Two of these hgcA-bins (from the Desulfobacterales order) were the second and third most abundant bins across our five metagenomes. The only two Geobacterales *hgcA* genes were binned, and no hgcA-bins were recovered from Geobacterales. We retrieved three hgcA+ Syntrophobacterales bins, one of which (SYN_0007) was the most abundant hgcA+ bin we recovered, accounting for nearly 3% of the overall bin coverage. We also recovered HgcA sequences that clustered with Clostridia-derived HgcA sequences. The main group of these sequences, including 4 binned HgcA sequences, forms a monophyletic cluster with weak bootstrap support (Figure S3). However, there were also two hgcA+ Clostridia bins (CLO_0015, CLO_0016) with HgcA sequences that fall outside of this monophyletic cluster and clustered weakly with an array of divergent sequences, highlighting the limitations of an exclusively assembly-based analysis as compared to genome-resolved binning. Other previously known methylators that were detected in our metagenomes included methanogenic archaea (4 HgcA sequences, 3.5% of *hgcA* coverage, 1 hgcA+ bin) and Chloroflexi (5 HgcA sequences, 1.3% of *hgcA* coverage, and 1 hgcA+ bin). Overall, methanogenic archaea were very rare, with only one bin accounting for about a half percent of the total bin coverage. Chloroflexi bins accounted for ∼3% of the total metagenomic reads, but most of these reads came from hgcA-bins in different orders than the hgcA+ bin.

The majority of *hgcA* read coverage was accounted for by two large groups of bacteria, neither of which are experimentally verified methylators. Fourteen of these sequences, accounting for 13% of the total coverage, formed a monophyletic cluster with substantial bootstrap support (Figure S3). Taxonomic analysis by GTDB and the rp16-based phylogeny of the four hgcA+ bins in this cluster identified them as Bacteroidetes. This is supported by the co-clustering of HgcA sequences from Bacteroidetes bins downloaded from NCBI, which are mostly bins reconstructed from aquifer metagenomes, but also include a bin from a thiocyanate reactor^53, 65–67^. The four hgcA+ bins from this study were all within a subset of the Bacteroidales order, which also contained eight hgcA-bins (Figure S7). An additional 12 hgcA-Bacteroidales bins fall outside of the above-mentioned cluster. We did identify one hgcA+ Bacteroidales isolate, *Paludibacter jiangxiensis*, that was cultivated from a rice paddy field,^68^ but the methylation phenotype has not been experimentally confirmed in this species or any other Bacteriodales member. The other large cluster of 33 HgcA sequences accounted for 50% of the total *hgcA* coverage. We could only recover a few genes from the NCBI non-redundant database that clustered with these sequences, and none from reference isolate genomes. Phylogenetic analysis of the 15 bins with these HgcA sequences identified them as members of the Planctomycetes-Verrucomicrobia-Chlamydia (PVC) superphylum (Figure S6, Figure 3). Bin rp16 phylogenies show that 11 of the bins are within the newly-proposed Kiritimatiellaeota phylum,^69^ two are from the Lentisphaerae phylum, and one each from the Verrucomicrobia and Planctomycetes phyla. The PVC superphylum dominates the overall read coverage of our bins as well, with 79 PVC bins accounting for 42% of total bin coverage. The Kiritimatiellaeota phylum alone accounts for 37 bins and ∼30% of total bin coverage, including four of the eight most abundant bins (Figure 3, Data File 1). There are very few publicly available Kiritimatiellaeota genomes and only one cultured representative.^69, 70^ Notably, a recent paper also identified several hgcA+ bins associated with the Kiritimatiellaeota phylum in a sulfate-enriched lake, but the HgcA sequences from those bins did not cluster closely to those from this study (Figure S3). Instead, they clustered just outside of the PVC superphylum cluster, along with two unbinned HgcA sequences from our metagenome contigs. This is consistent with the rp16-based bin phylogeny, where Kiritimatiellaeota genomes from Jones et al., (2019) were also distinct from the Mendota hgcA+ Kiritimatiellaeota (Figure 3). For both the Kiritimatiellaeota and the Bacteroidales, the presence of *hgcA* within bins was not phylogenetically conserved (Figure 3, Figure S7). That is, bins with and without *hgcA* genes cluster together within these lineages. Additionally, the hgcA+ bin read coverage from these two groups was not significantly different from their hgcA-counterparts (data not shown). We also identified several other novel putative methylators that were lower in number and abundance, including Margulisbacteria, Firestonebacteria, and Actinobacteria.

**Figure 3.**
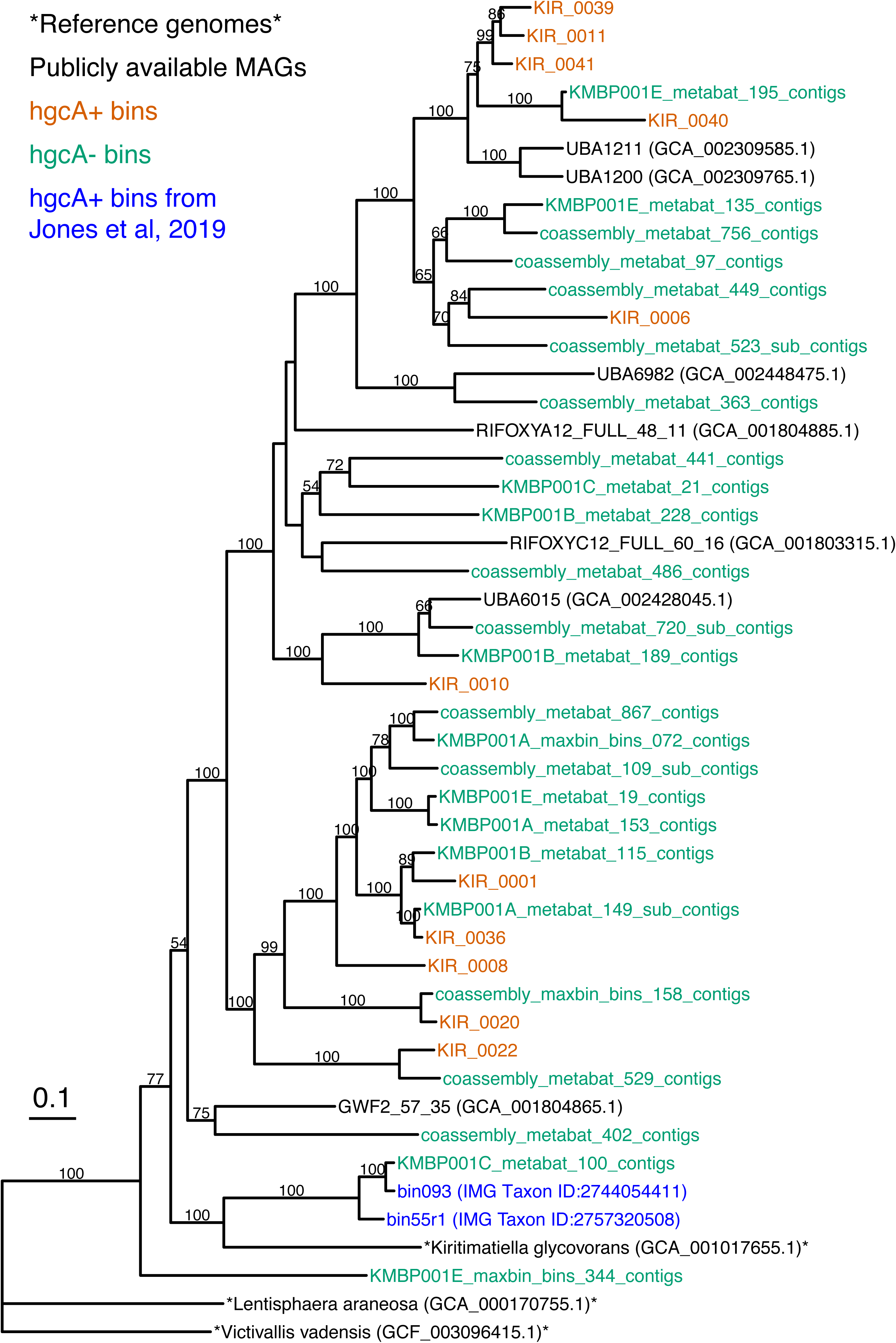
The hgcA gene is widespread in Mendota Kiritimatiellaeota bins, but is not phylogenetically conserved. Maximum-likelihood tree is based on a concatenated alignment of rp16 proteins. Names in orange are hgcA+ bins from this study, and green names are hgcA-bins from this study. Names in black are genomes or bins pulled from NCBI, and genomes with the asterisks indicate cultured isolate reference genomes. The accession version numbers are in parentheses following the bin or genome name. The bin names in blue correspond to two hgcA+ bins from a recent publication.^1^ The tree was generated in RAxML and rooted using the two Lentisphaerae genomes (*Lentisphaera araneosa* and *Victivallis vadensis*). Bootstrap values of less than 50 are not shown.

**Figure 4.**
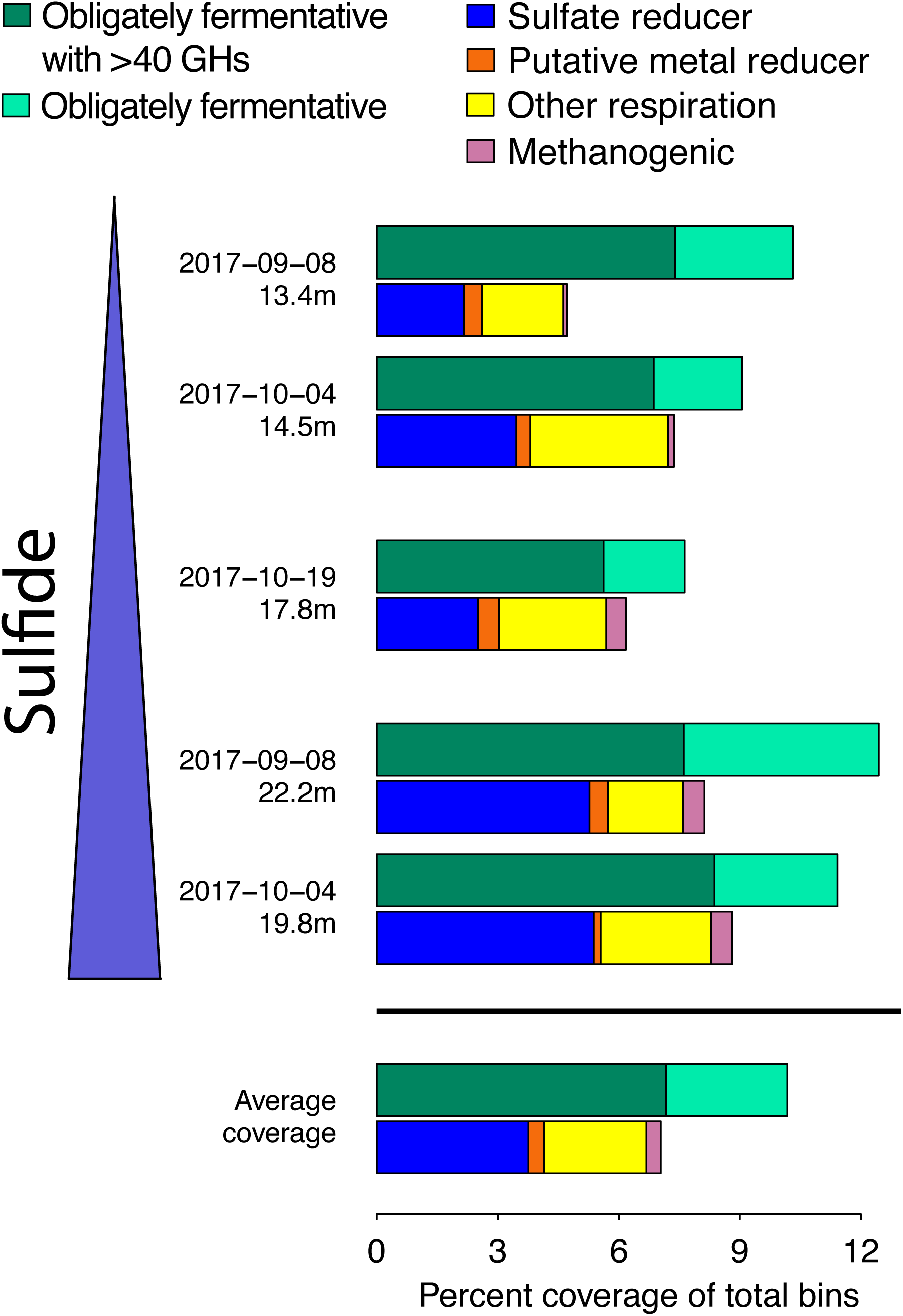
Fermentative organisms are the most abundant hgcA+ organisms in Lake Mendota. Total coverage of hgcA+ bins in different metabolic guilds across each metagenome. Plots of coverage in the different metagenomes are arranged by decreasing redox potential, which corresponds to increasing sulfide concentrations. Abbreviations: GHs - glucoside hydrolases.

### Metabolic potential of methylating bins

To understand how methylating organisms in Lake Mendota might be linking biogeochemical processes to MeHg production, we explored the major metabolic pathways encoded in our hgcA+ bins. Due to the abundance of sulfate in Lake Mendota and the extensive literature linking MeHg production to SRBs, we hypothesized that most hgcA+ genomes would harbor genes enabling sulfate reduction. Three of the four Desulfobacterales genomes contained the *dsrAB* gene cluster, *dsrD*, *aprAB*, *sat*, and the *qmoABC* genes requisite for sulfate reduction, with only DES_0017 lacking these genes (Figure S8). SYN_0007, the highest coverage hgcA+ bin, and SYN_0037 also contained this set of canonical sulfate-reduction genes. Just over half of the SRBs by coverage (52%) were hgcA+, and there were only three hgcA-sulfate reducers. Five of our hgcA+ bins, including two of the aforementioned sulfate reducers, contain molybdopterin oxidoreductase (MoOR) sequences that are homologous to polysulfide reductase (*psr*) (Figure S9). These genes are in neighborhoods with the classic complex iron–sulfur molybdoenzyme (CISM) architecture, with downstream Fe-S binding proteins and an integral membrane anchor protein similar to the nrfD protein (Figure S9).^71^ Some of these clusters also have rhodanese domain-containing proteins nearby. These genes likely confer the ability to respire partially reduced inorganic sulfur compounds such as tetrathionate or thiosulfate.^71^ However, these three bins also have the genetic machinery to mediate other terminal respiration processes and are unlikely to be exclusively reliant on the psrA for respiration.

MeHg production has also been linked to Fe reduction. We retrieved several hgcA+ bins with potential for extracellular electron transfer (EET), which is often used to respire insoluble metal complexes such as Fe or Mn oxides (Figure S10). Both Geobacterales bins have a porin-cytochrome c complex (PCC) operon that is homologous to the ExtEFG operon from *Geobacter sulfurreducens* and 23 and 25 MHC proteins, respectively (Figure S8, S10).^72^ The two Geobacterales bins had very low read coverage, but were most abundant in CHE3, where we saw evidence for enhanced Mn cycling and peaks in fraction MeHg (Data File 1). Combined with the observation that Geobacterales methylators often produce MeHg at a high rate in culture, this suggests that Mn cycling at the oxic-anoxic interface may be playing a role in MeHg production. To our knowledge, Mn reduction has not previously been associated with Hg methylation. However, further work is needed to experimentally verify this. These are the only two bins from this study with PCCs homologous to experimentally verified metal-reducing complexes. The PCC operon identified in VER_0023 is homologous to PCCs identified in Verrucomicrobia genomes recovered from bog lakes, where they are thought to mediate the dispersal of electrons onto either iron or humic substances. However, to our knowledge the function of these PCCs have not been experimentally verified.^73^ The other PCC operons were found in Bacteroidetes and Kiritimatiellaeota bins and were not closely related to gene clusters with experimentally verified EET function, but the corresponding organisms appear to be capable of respiration (complex I, complete TCA cycle). The relatively low coverage of these bins at depths with enhanced Mn cycling suggests that EET-mediated Mn reduction is not their primary respiratory pathway (Data File 1).

We also detected the machinery for nitrogen species reduction in our methylating organisms (Figure S8). GEO_0030 and DES_0034 have at least one nitrate reductase (*napDAGHB*, *narG*) and *nrfA*, suggesting they are capable of mediating dissimilatory nitrate reduction to ammonia (DNRA). Three other hgcA+ bins (PLA_0021, KIR_0036, DES_0019) have only the *nrfHA* gene cluster, and thus likely support nitrite reduction to ammonia. While *nrfHA* can be used for energy conservation, it is also implicated in nitrite detoxification, sulfite reduction, and reducing power dispersal during fermentation.^74, 75^ This, in combination with low nitrate/nitrite levels in the water column during this time of year (data not shown) and the presence of sulfur cycling genes and/or PCCs in these bins suggest that nitrogen-based respiration does not play a major role in MeHg production in this system. The various DNRA and denitrifying genes, as well as oxidases, were wide-spread throughout the hgcA-bins we recovered. However, due to the lack of oxidized nitrogen species and oxygen throughout the hypolimnion during this time, we suspect that these bins correspond to either facultative aerobes that are living fermentatively or that are metabolically inactive at these sites.

We only recovered one bin derived from a methanogen (MET_0028) that accounted for 0.3% of the total read coverage and was hgcA+ (Figure S7). This bin was most abundant in the highly reduced deep hypolimnetic samples. MET_0028 is a member of the Methanomicrobiales order, the current representatives of which are strictly CO_2_-reducing methanogens, using either formate or H_2_ for reducing power. Indeed, MET_0028 has a number of hydrogenases, formate dehydrogenase, and a complete Wood-Ljundahl pathway. The corresponding methanogen is likely dependent on the production of H_2_ and/or formate by fermentative organisms. The overall scarcity of methanogens suggests that methanogenesis in the water column is insignificant, and that sulfate reduction is the driving terminal respiratory process in the anoxic hypolimnion.

The remaining 27 hgcA+ bins lack canonical machinery for terminal electron-accepting processes and are likely to be involved in fermentative or syntrophic lifestyles (Figure S11). Several of these bins do have *cyd* operons, encoding cytochrome bd oxidases, but the lack of other machinery facilitating a respiratory metabolism suggests that these are used to minimize oxidative stress.^76^ Some bins also have the *nrfHA* operon, but likely use this for nitrite detoxification or fermentative metabolism.^74, 75^ One likely mechanism for electron dispersal among these groups is through hydrogen production, since 24 of them have hydrogenases commonly involved in the fermentative evolution of H_2_, mostly [FeFe] Group A hydrogenases, with some from [NiFe] Group 4. Fourteen of these, including all of the highly abundant Kiritimatiellaeota hgcA+ bins, also have an Rnf complex, which can facilitate reverse electron transport to drive the replenishment of oxidized ferredoxin and NAD+ when coupled with the evolution of H_2_ from confurcating hydrogenases such as those found in these genomes.^77^

Coupled with the prevalence of hydrogenases enabling H_2_ consumption in bins carrying respiratory machinery (both hgcA+ and hgcA-), these data hint that H_2_-mediated syntrophy could be an important driver of both overall community metabolism and MeHg production in Lake Mendota. We also looked at pathways potentially conferring the ability to ferment via pyruvate. Nearly all of the hgcA+ bins likely corresponding to fermenters had pyruvate:ferredoxin (PFOR), and many of them also had acetate kinase (*ack*) and phosphate acetyltransferase (*pta*), which together can mediate pyruvate fermentation to acetate. Eight hgcA+ bins had pyruvate formate lyase (pflB). This mediates another pyruvate fermentation pathway, where pyruvate is broken down to acetyl-CoA and formate. The formate can either be taken up for biosynthetic purposes through formate dehydrogenase (FDH) or exported and used by other organisms as an electron donor. While none of the hgcA+ bins corresponding to obligate fermenters encode FDH, all of the sulfate-reducer bins and many more of the bins carrying other respiratory machinery do have FDH, suggesting that the formate formed is exported and used by other organisms. Notably, the eight pflB+ bins fall into two distinct phylogenetic clusters, one within Kiritimatiellaeota and one within Clostridia (data not shown). They also possess an array of aldehyde and alcohol dehydrogenases that could facilitate the production of a range of short chain fatty acids. Bins linked to obligate fermentation were also common in the total microbial community, as they represent 106 of the 228 bins, accounting for almost 50% of the bin coverage. This does not include the many bins containing genes for dissimilatory nitrogen reduction or oxidases that were likely maintaining fermentative metabolism at these depths. Many of the hgcA+ bins associated with obligate fermentation also encoded genes mediating the primary degradation of particulate organic carbon. Thirteen of these bins have at least 40 glucoside hydrolases (GHs), suggesting that their associated organisms are adapted to degrading polysaccharides. The highly abundant Kiritimatiellaeota hgcA+ bins, in particular, appear to be well adapted to polymer degradation, with bins carrying up to 468 GHs. In fact, 100 total bins each carried over 40 GH genes, suggesting that primary polysaccharide degradation is one of the dominant metabolic strategies in the anoxic water column in Lake Mendota. Of these, 49 represent obligate fermenters, while 50 are suspected to represent facultative aerobes. One bin with 44 GHs corresponds to a sulfate-reducing Chloroflexi. The bins also possess a wide diversity of peptidases, although these genes are also prevalent among the hgcA+ bins associated with respiratory lifestyles.

We know little about the mass flux constraints on carbon degradation in the anoxic water column of freshwater lakes, and even less about how they influence the production of MeHg. In other natural anoxic environments such as marine sediments, the hydrolysis and primary fermentation of large polymers, such as polysaccharides, is thought to be the rate-limiting step in anoxic microbial community metabolism.^78–80^ Lake Mendota is a eutrophic system with a multi-year residence time, and thus the DOC pool is dominated by autochthonous inputs.^81^ Primary production in the lake is dominated by cyanobacteria, which have a high proportion of EPS in their biomass.^82, 83^ This could explain the abundance of polysaccharide-degrading obligate fermenters, which account for ∼50% of both the hgcA+ and overall bin coverage. Such organisms break down and ferment carbohydrates, producing smaller carbon compounds that can be further metabolized by secondary fermenters or syntrophs, or directly consumed by respiratory bacteria. Of the bins linked to respiration, SRB accounted for 22% of the total hgcA+ bin coverage and only 7% of the overall bin coverage. Methanogenic archaea and EET-capable Geobacterales bins accounted for a very small percentage of both the hgcA+ and the overall read coverage. Aerobic and nitrate/nitrite-reducing organisms were also prevalent, but due to the redox status of the sampling depth we do not expect them to be respiring at these depths. Thus, sulfate reduction is likely the dominant respiratory pathway in this community, despite the relatively low abundance of SRBs. The breakdown of large polymers and their subsequent degradation through the anaerobic food web drive community metabolism, and probably also drive MeHg production. However, we know very little about differences in MeHg production between different hgcA+ microbes or the specific rate-limiting steps for microbial community metabolism in this system. This highlights the need for further research to probe how biogeochemical conditions can indirectly influence MeHg production *in situ* by mediating changes in the carbon and energy flux through the anaerobic food web.

## Acknowledgements

The authors acknowledge the North Temperate Lakes Long Term Ecological Research (NTL-LTER) site, Lake Mendota Microbial Observatory field crews, UW Center for Limnology for field and logistical support. In particular we thank graduate students Tylor Rosera, Stephanie Berg, and Marissa Kneer for sampling assistance, and Joseph Skarlupka for computational assistance. We also thank undergraduate researchers Mykala Sobieck for sampling design and sampling assistance, and Diana Mendez and Ariel Sorg for sampling assistance. Geochemical analyses were performed at the Water Science and Engineering Laboratory at the University of Wisconsin – Madison. Mercury analyses were performed in the Mercury Research Laboratory in the Water Science Center at the U.S. Geological Survey in Middleton, WI. This research was performed in part using the Wisconsin Energy Institute computing cluster, which is supported by the Great Lakes Bioenergy Research Center as part of the U.S. Department of Energy Office of Science. Katherine D. McMahon received funding from the United States National Science Foundation Microbial Observatories program (MCB-0702395), the Long-Term Ecological Research Program (NTL–LTER DEB-1440297), and an INSPIRE award (DEB-1344254), the Wisconsin Alumni Research Foundation, and the National Oceanic and Atmospheric Administration (NA10OAR4170070 via the Wisconsin Sea Grant College Program Project #HCE-22). Benjamin Peterson was supported by the National Science Foundation Graduate Research Fellowship Program under grant no. XXXXX during this research.

## Supplementary Figures

**Figure S1.**
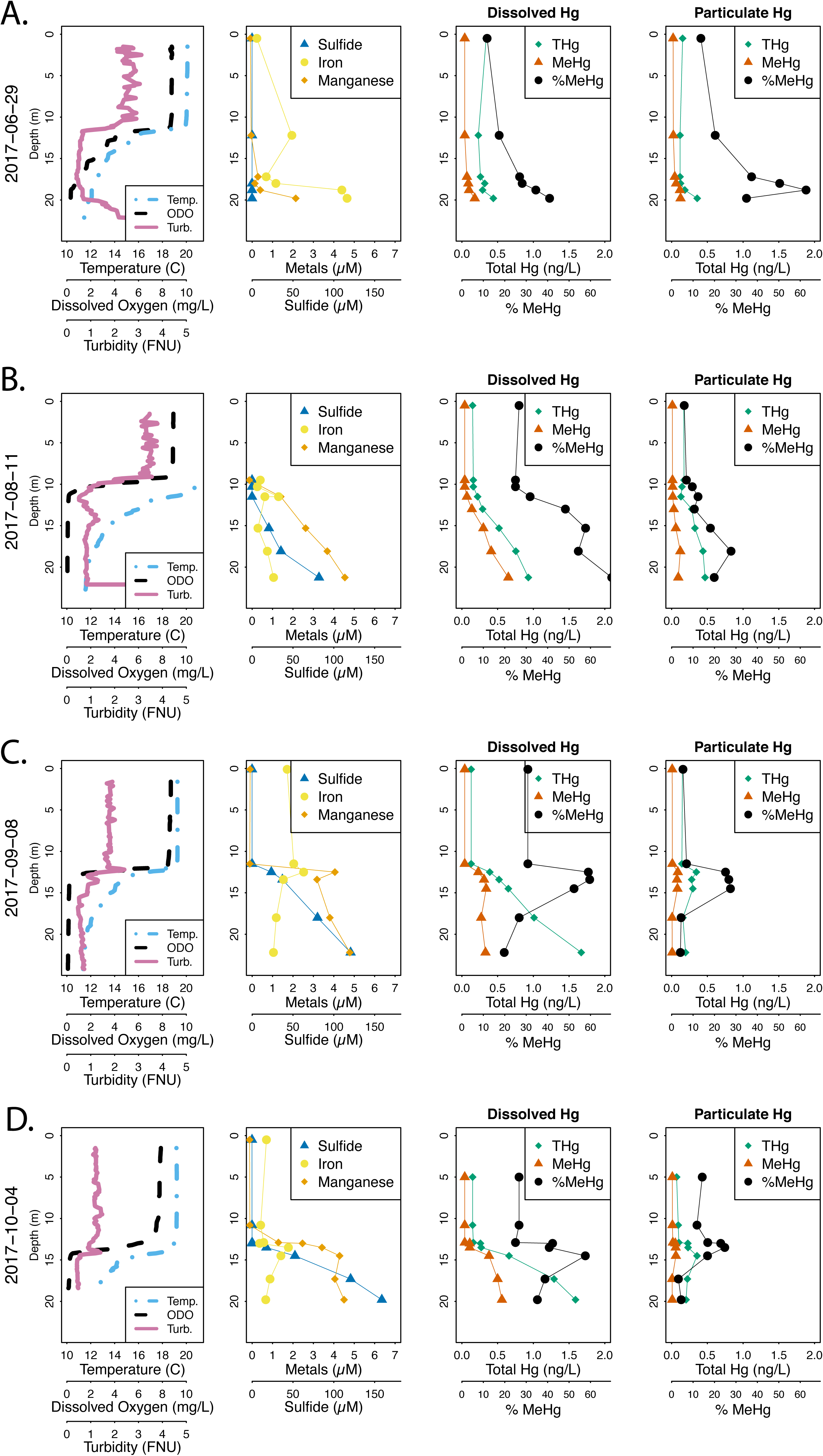
Representative profiles of Lake Mendota from across the open water season in 2017. The dissolved Hg species are operationally defined as everything that passes a quartz fiber filter (QFF), and the particulate fraction is what is retained on a QFF. Both iron and manganese are the dissolved fraction only (0.45µm PES filter). Abbreviations: Temp.-Temperature (°C), ODO - Optical dissolved oxygen in mg/L, Turb. - Turbidity in Formazin Nephelometric Units (FNU), THg - Total mercury, MeHg - Methylmercury, %MeHg - Methylmercury concentration divided by total mercury concentration.

**Figure S2.**
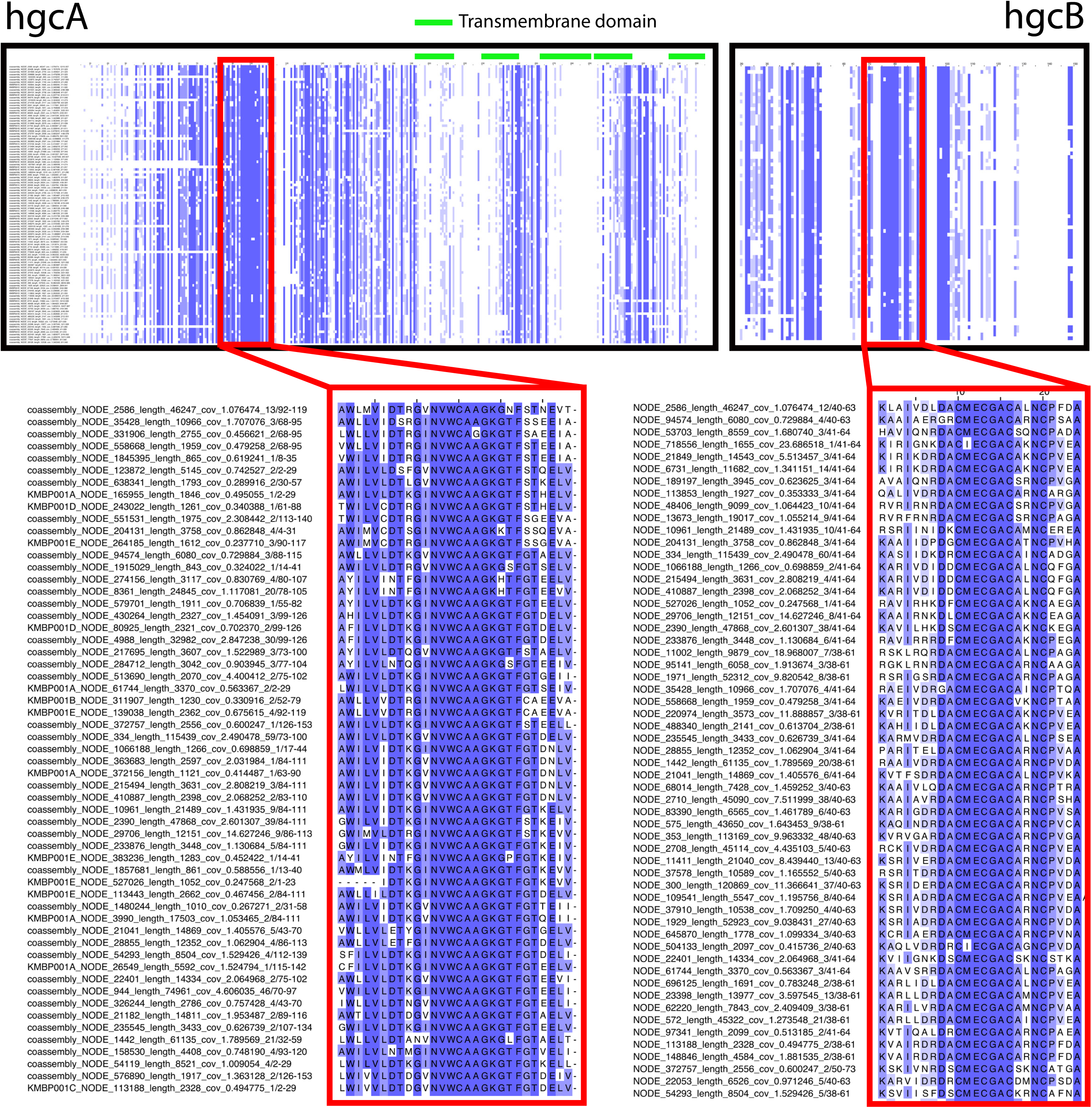
Alignments of identified hgcA and hgcB amino acid sequences from all five metagenomes. Green bars indicate regions of predicted transmembrane domains in the alignments. The zoomed-in portion of the hgcA alignment highlights a portion of the corrinoid-binding domain for a subset of the sequences, and includes the characteristic highly conserved cap-helix domain. For hgcB, we highlighted a portion of the alignment that includes one of the two highly conserved ferredoxin-binding motifs from a subset of the sequences.

**Figure S3.**
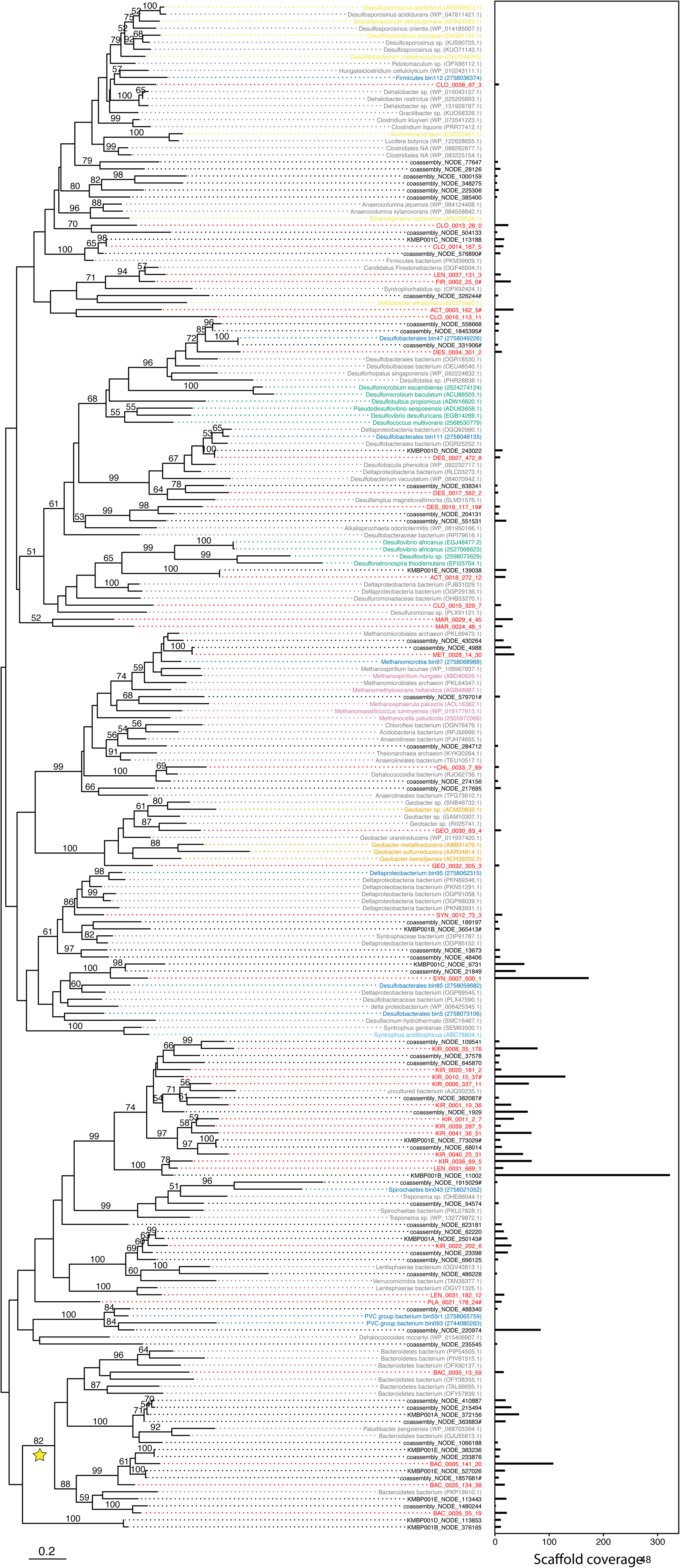
Maximum likelihood tree of hgcA sequences and overall coverage across five metagenomes. Names in black indicate unbinned hgcA sequences. For hgcA sequences that were binned, the scaffold name was replaced with the bin name (red names). Dark blue names indicate hgcA sequences from bins from a recent paper in a similar system.1 These bins are followed by the IMG Taxon ID in parentheses. Grey names indicate hgcA sequences downloaded from NCBI’s non-redundant database that did not come from the genome of a confirmed methylating organisms. Remaining colored names are from genomes of confirmed methylators and match the color scheme in Figure 2 (yellow - Clostridia; green - Desulfobacterales; pink - methanogens; orange - Geobacterales; light blue - Syntrophobacterales). All reference sequence names are followed by their accession version number in parentheses. Scaffold coverage is the average coverage of nucleotides in the corresponding hgcA+ scaffold across all five metagenomes. Sequence names from this study that are followed by a pound sign do not have a trailing hgcB sequence.

**Figure S4.**
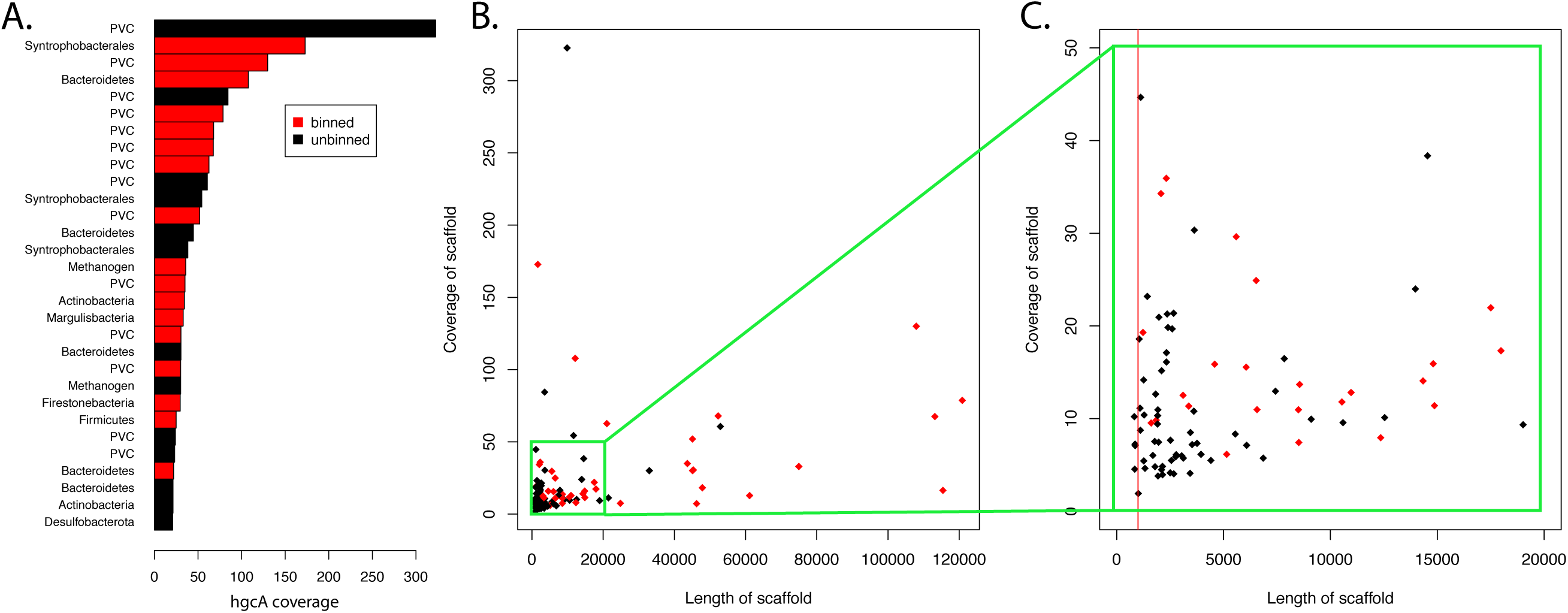
Overview of binning of hgcA sequences. A: Rank abundance curve of hgcA sequences across all metagenomes. Bars colored in red indicate a binned sequence. B: Plot of average coverage of scaffold vs. length of scaffold of hgcA sequences, with red dots indicating that the sequence was binned.

**Figure S5.**
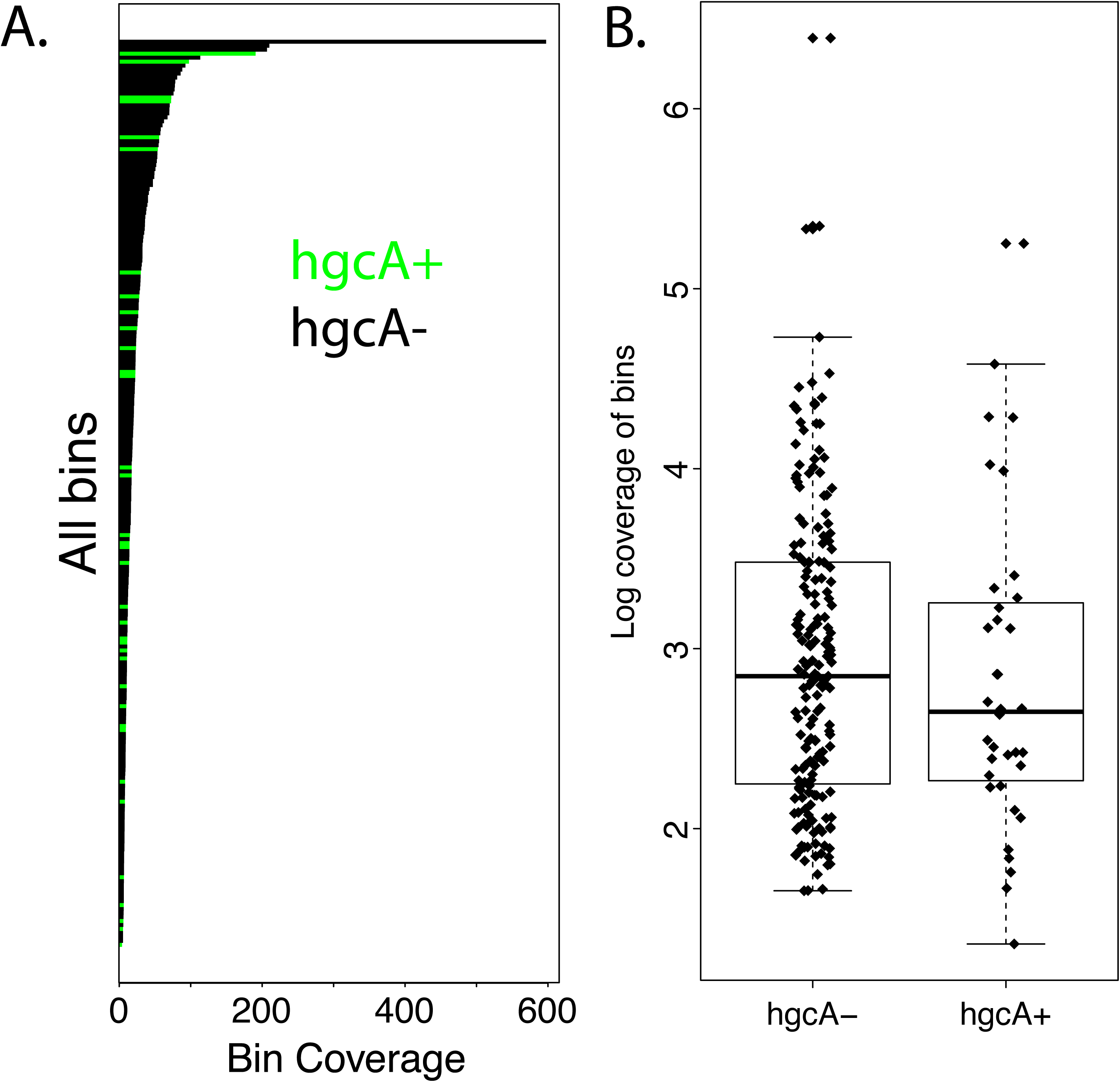
Comparison of coverage between hgcA+ and hgcA- bins. A: Rank abundance curve of all bins across all metagenomes. Bins encoding hgcA are colored green. B: Log coverage of hgcA+ vs. hgcA- bins.

**Figure S6.**
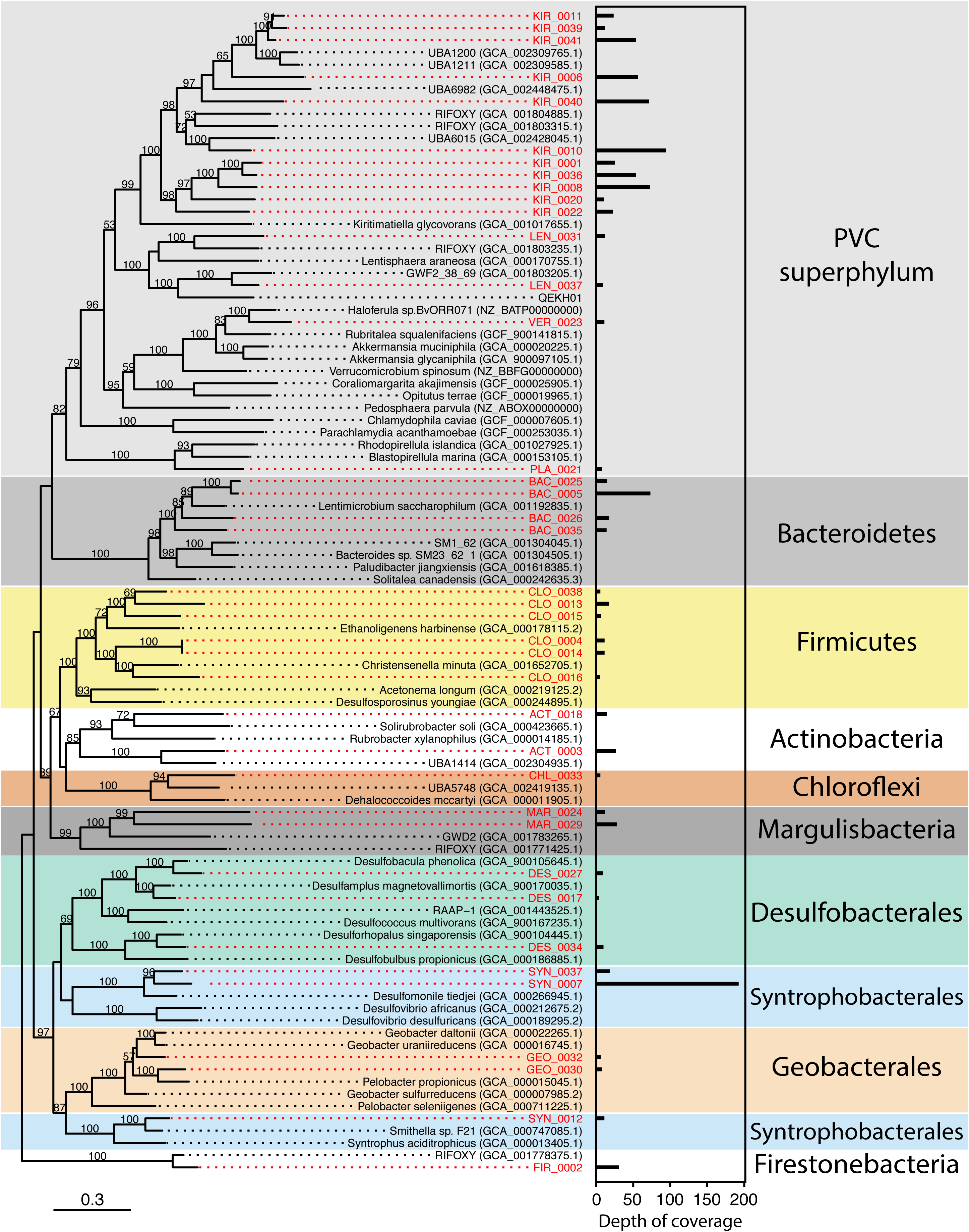
Maximum likelihood tree of rp16 genes from all bacterial hgcA+ bins and reference genomes from NCBI. Bootstrap values below 50 have been removed. Tree was rooted using three archaeal bins from this study.

**Figure S7.**
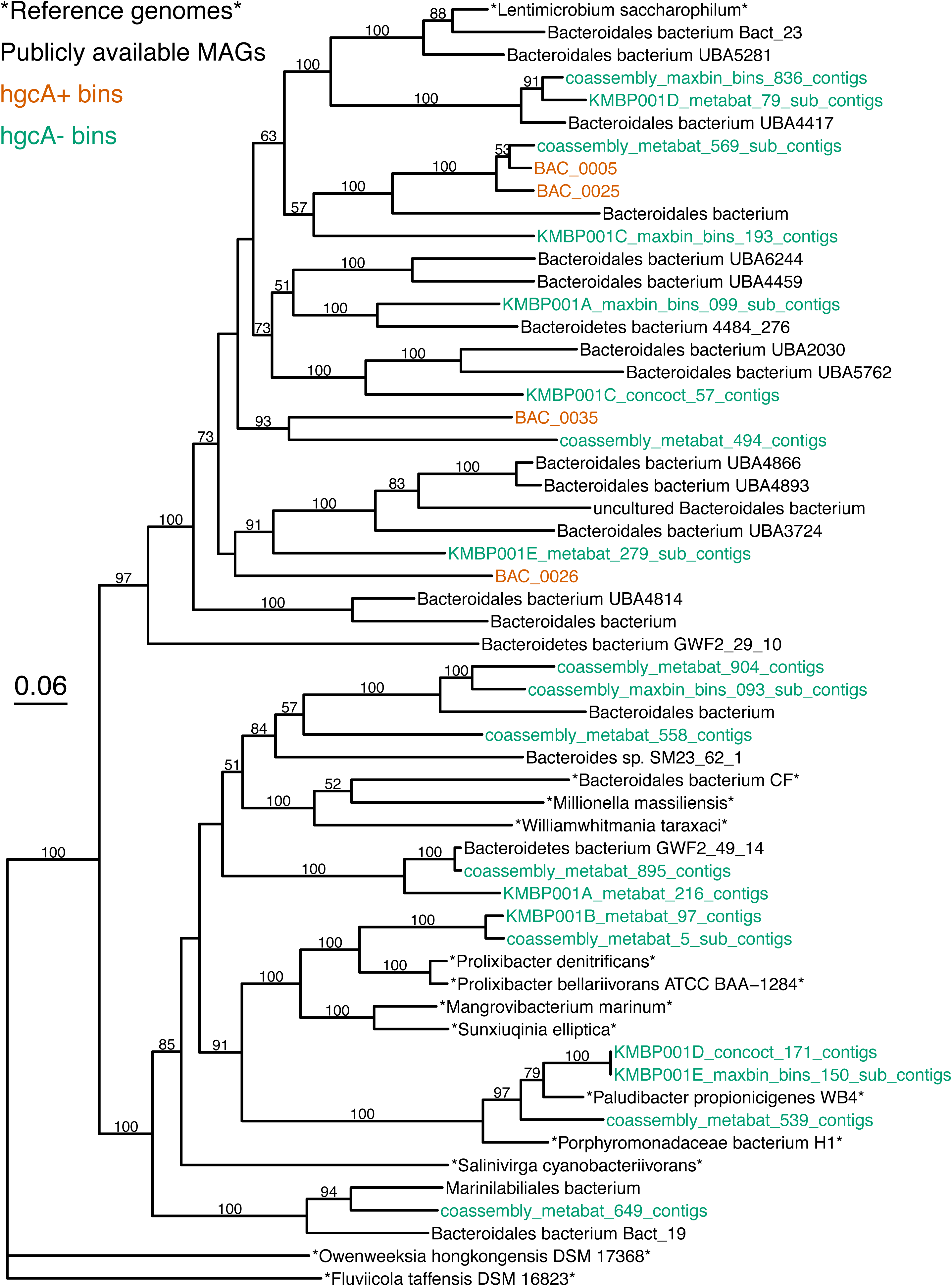
Maximum likelihood tree of rp16 gene from all Bacteroidales bins from this study. Bin names in green are hgcA- bins, while those in orange are hgcA+ bins. Sequences in black are bins downloaded from NCBI, and bin names surrounded by asterisks are reference genomes from isolate cultures. The Bacteroidales tree was rooted using two Flavobacteriales reference genomes (Owenweeksia hongkongensis DSM 17368 and Fluviicola taffensis DSM 16823).

**Figure S8.**
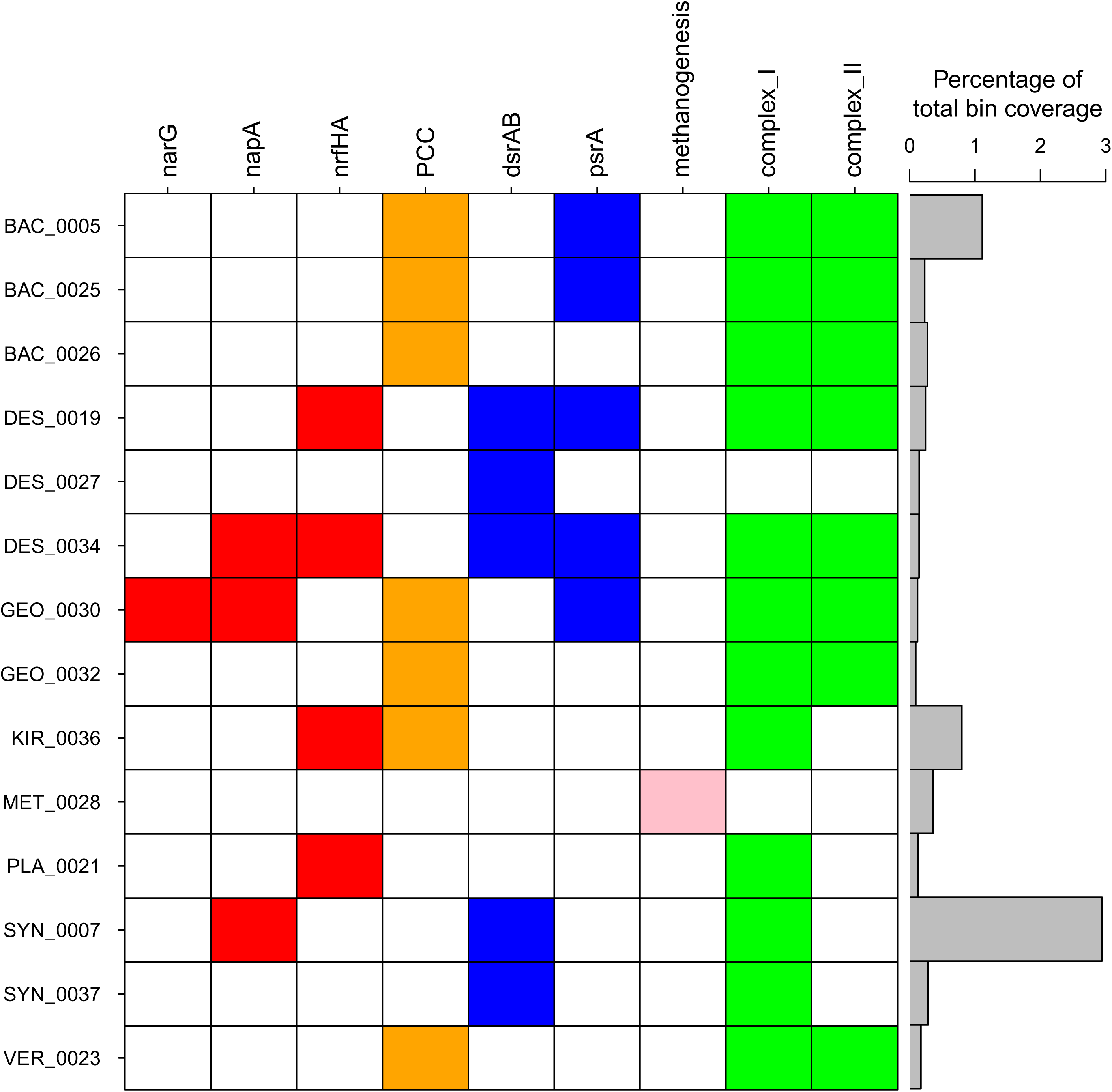
Heatmap of metabolic potential of hgcA+ bins with respiratory metabolic genes and overall bin abundance. Dissimilatory nitrogen cycling genes are in red: narG = membrane-bound nitrate reductase, napA = periplasmic nitrate reductase, nrfHA = cytochrome c nitrite reductase. Genes for nitrite reduction by denitrification were not identified in any hgcA+ bins. Putative external electron transfer proteins are in orange: PCC = Porin-cytochrome c complex. Sulfur cycling genes in blue: dsrAB = dissimilatory sulfite reductase; psrA = polysulfide-reductase homolog. Methanogenesis refers to the overall phenotype indicated by the bin. In green are complex I (the 11 and/or 14 subunit version) and complex II of the electron transport chain.

**Figure S9.**
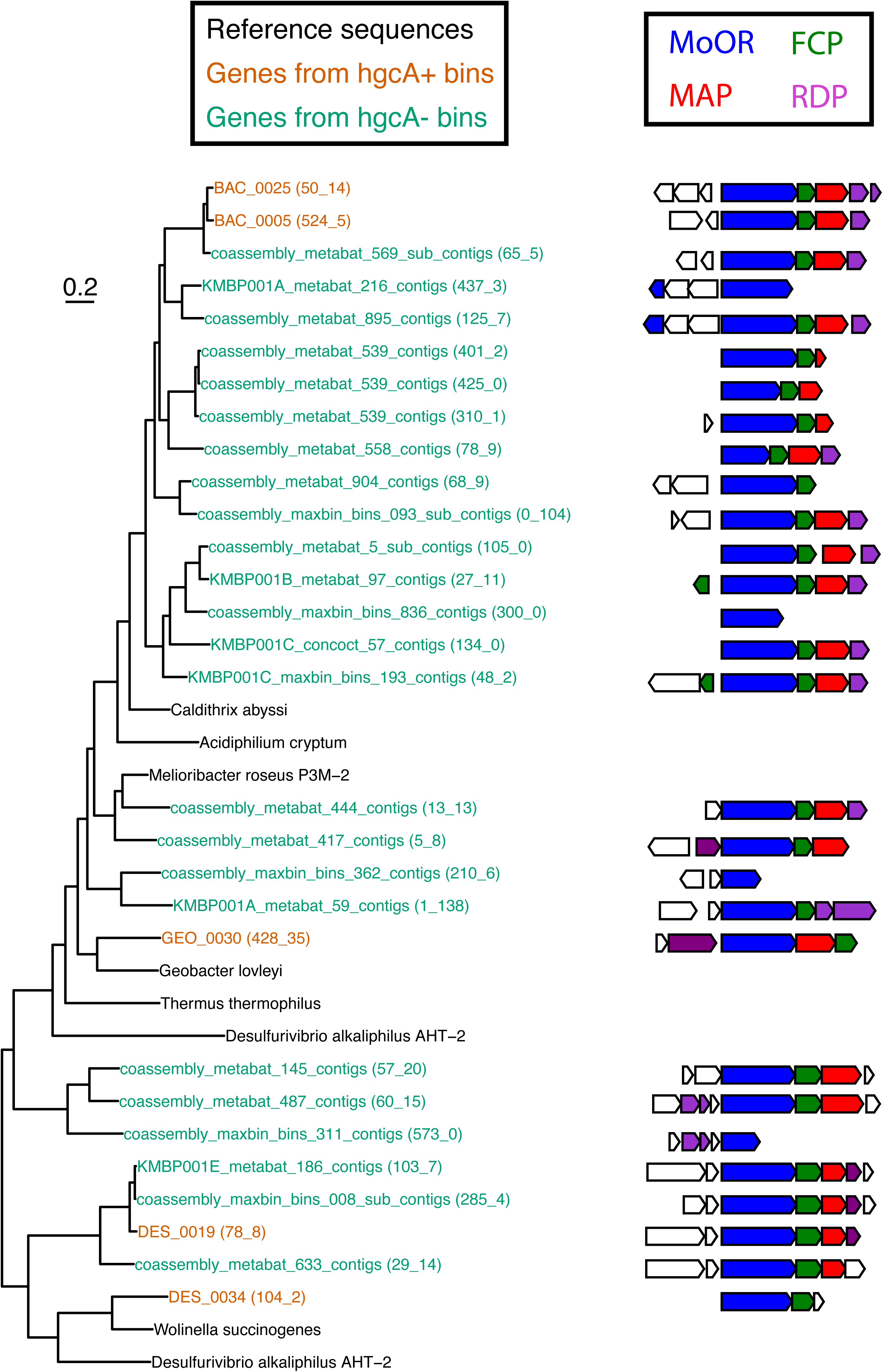
Phylogenetic tree of polysulfide reductase (psr) homologs from hgcA+ bins. In the branch labels, the bin names are followed by the scaffold number and ORF number in parentheses. Names in orange are from hgcA+ bins, green are from hgcA-bins. Names in black correspond to reference sequences. The gene neighborhoods within 2500bp upstream and downstream of the corresponding MoOR from this study are shown to the right of the tree. The canonical complex iron–sulfur molybdoenzyme (CISM) architecture includes the MoOR (shown in blue), a four-cluster protein (FCP) with four Fe-S clusters (shown in green), and a membrane anchor protein (MAP), such as the nrfD subunit from the nitrite reductase complex NrfABCD (shown in red). Rhodenase-domain proteins (RDP), involved in sulfur transport, are shown in purple.

**Figure S10.**
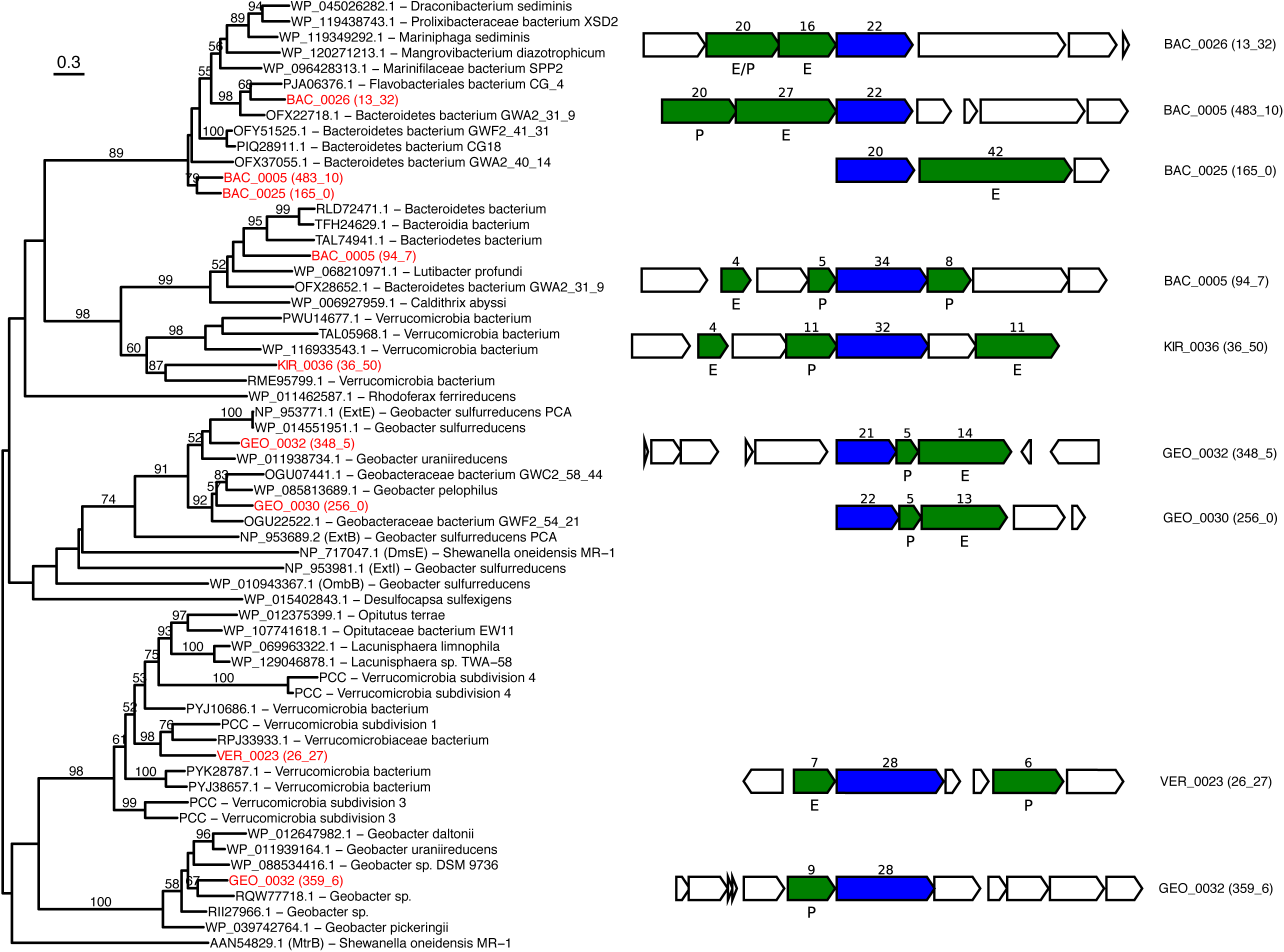
Phylogenetic tree and gene neighborhoods of beta-barrel outer membrane protein (BB-OMP) genes from hgcA+ bins. Sequence names in red are from hgcA+ bins, and the following numbers in parentheses indicate the scaffold and ORF, respectively. The gene neighborhoods within 4000bp upstream and downstream of the BB-OMP genes are shown to the right of the tree. BB-OMP sequences are shown in blue, and the predicted number of transmembrane sheets within the protein are provided above the gene. Predicted multiheme cytochrome c proteins are shown in green, with the number of the heme-binding sites above the gene. The predicted localization of the protein is shown below the gene (E indicates extracellular, P indicates perisplasmic).

**Figure S11.**
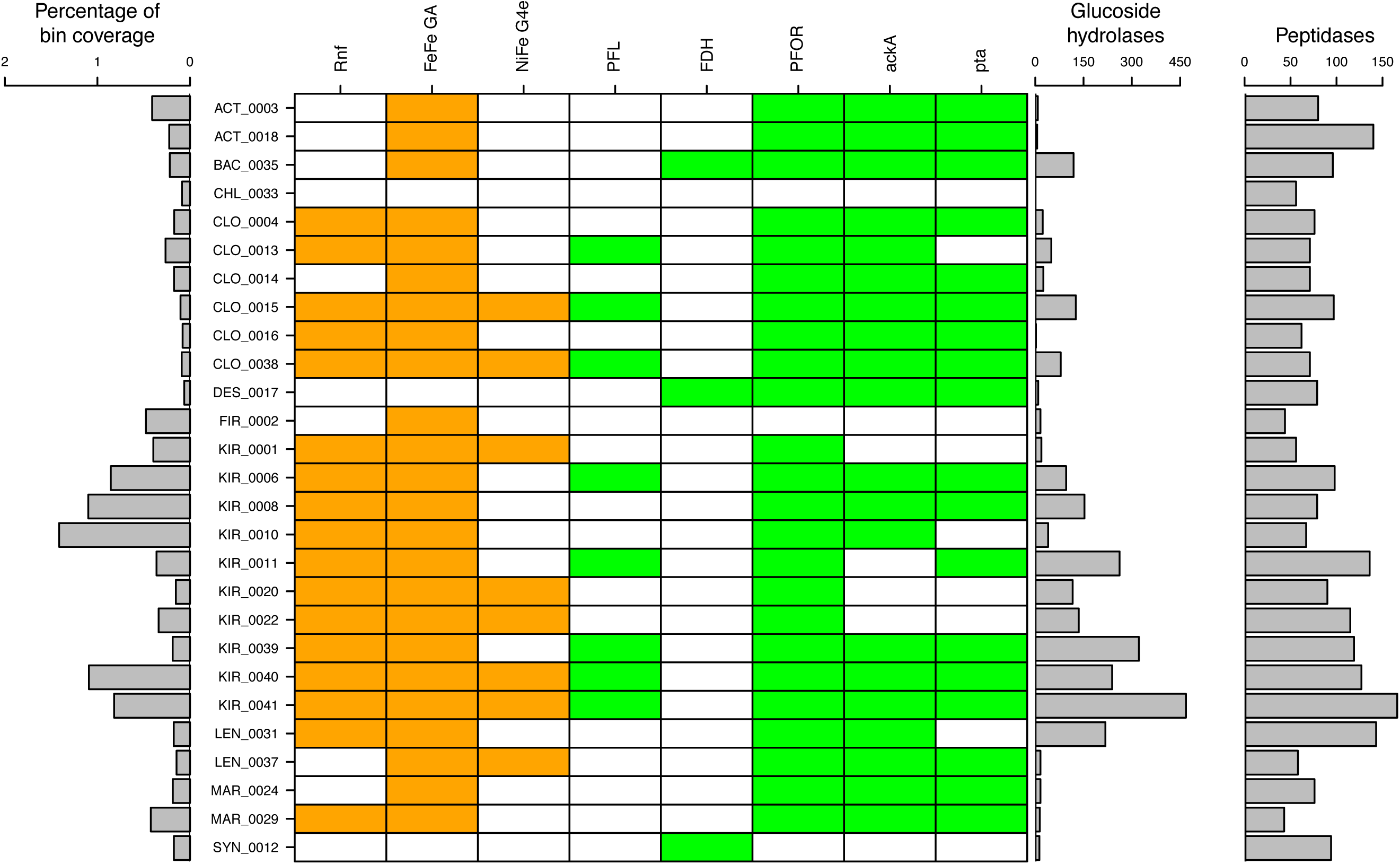
Abundance and metabolic gene features of fermentative bins. The percentage of bin coverage is relative to the total coverage of all the bins from this study. Genes potentially involved in fermentative hydrogen evolution are shown in orange: Rnf = Rhodobacter nitrogen fixation complex; FeFe GA = [FeFe]-hydrogenase, group A; NiFe G4e = [NiFe]-hydrogenase, group 4e. Genes or gene clusters involved in fermentation of pyruvate are shown in green: PFL = pyruvate-formate lyase; FDH = formate dehydrogenase; PFOR = pyruvate-ferredoxin oxidoreductase; ackA = acetate kinase (ADP-forming); pta = phosphate acetyltransferase.

**Figure S12.**
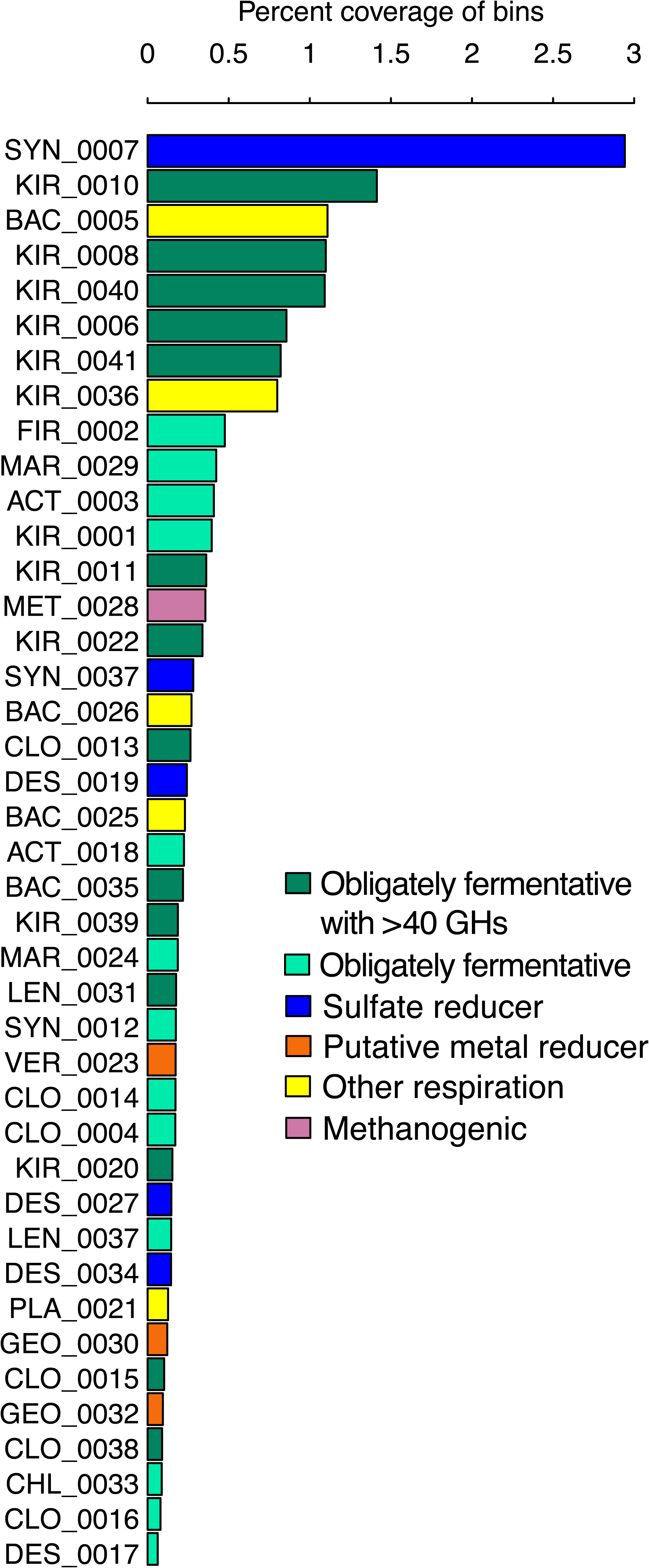
Rank abundance curve of hgcA+ bins across all five metagenomes, colored by predicted metabolic potential. The bin coverage is relativized to the total coverage of all the bins (both hgcA+ and hgcA-).

**Table S1.**
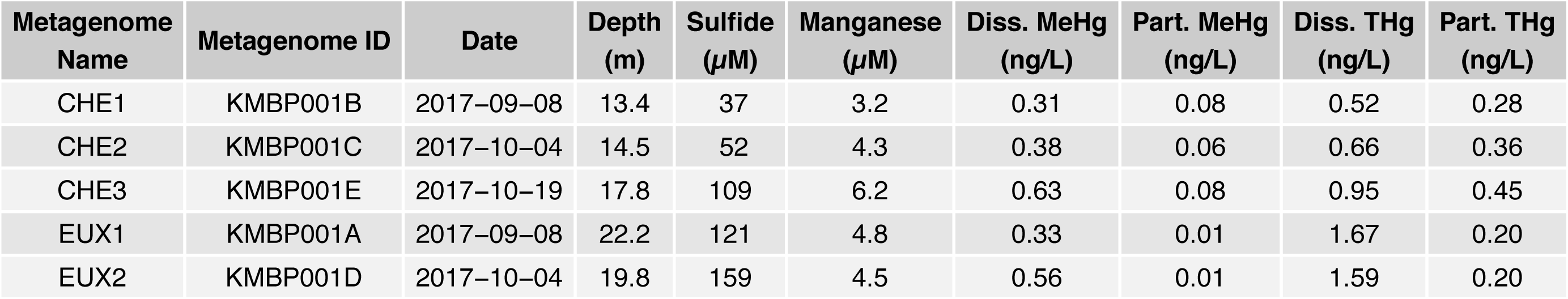
Summary of metadata and geochemical data associated with metagenomic samples.

**Table S2.** Assembly statistics for each of the assemblies, after removing all scaffolds <500bp in length.

| Assembly ID | Number of scaffolds | N50 | L50 | Total assembly length (bp) | Total number of ORFs |
| --- | --- | --- | --- | --- | --- |
| CHE1 | 1,185,495 | 1630 | 201,484 | 1.603e+09 | 4,160,915 |
| CHE2 | 1,241,217 | 1528 | 227,330 | 1.622e+09 | 4,254,833 |
| CHE3 | 1,613,022 | 1471 | 306,579 | 2.057e+09 | 5,639,867 |
| EUX1 | 1,319,783 | 1486 | 235,323 | 1.698e+09 | 4,641,626 |
| EUX2 | 1,111,274 | 1307 | 227,854 | 1.327e+09 | 3,862,047 |
| coassembly | 3,908,837 | 1844 | 611,093 | 5.651e+09 | 13,999,347 |

## Data File Legends.

**Data File 1.** Bin information and statistics. Completeness and redundancy estimates are based on universal conserved proteins set in CheckM. Inferred taxonomy is based on a rp16-based ML-tree tree with a large reference data set. Coverage of each bin in each metagenome has been normalized to the number of reads in the smallest metagenome.

**Data File 2.** Custom hgcA Hidden Markov Model, based on HgcA amino acid sequences from 30 confirmed methylating isolates. See Supplementary Methods for details of HMM construction.

**Data File 3.** Fasta file of dereplicated amino acid sequences of HgcA sequences identified in assemblies.

**Data File 4.** Fasta file of dereplicated nucleic acid sequences of hgcA genes identified in assemblies.

**Data File 5.** Aggregated information for each hgcA gene from the dereplicated set. Classification of hgcA is based on the bin phylogenies, for the binned genes, and on the HgcA phylogenies, for the unbinned genes. “Rogue taxa” indicates that the HgcA sequence was highly divergent and interfering with phylogenetic reconstruction. These sequences were classified using pplacer with the hgcA phylogeny. The hgcB column indicates whether or not there was an hgcB gene immediately downstream of the hgcA gene on the scaffold. The abundance of each sequence is presented as the percentage of hgcA coverage within a metagenome that each gene accounts for.

## References

1. Amos HM, Jacob DJ, Streets DG, Sunderland EM. Legacy impacts of all-time anthropogenic emissions on the global mercury cycle. Global Biogeochem Cycles. 2013;27(2):410–421. doi:10.1002/gbc.20040

2. UN. Global Mercury Assessment.; 2018.

3. Bloom NS. On the chemical form of mercury in edible fish and marine invertebrate tissue. Can J Fish Aquat Sci. 1992;49(5):1010–1017. doi:10.1139/f92-113

4. Cleckner LB, Gilmour CC, Hurley JP, Krabbenhoft DP. Mercury methylation in periphyton of the Florida Everglades. Limnol Oceanogr. 1999;44(7):1815–1825. doi:10.4319/lo.1999.44.7.1815

5. Jensen S, Jernelöv A. Biological methylation of mercury in aquatic organisms. Nature. 1969;223:753–754. doi:10.1038/223754b0

6. Rothenberg SE, Feng X. Mercury cycling in a flooded rice paddy. J Geophys Res. 2012;117(G3003). doi:10.1029/2011JG001800

7. Sunderland EM, Krabbenhoft DP, Moreau JW, Strode SA, Landing WM. Mercury sources, distribution, and bioavailability in the North Pacific Ocean: Insights from data and models. Global Biogeochem Cycles. 2009;23(2). doi:10.1029/2008GB003425

8. Watras CJ, Bloom NS, Claas SA, Morrison KA, Gilmour CC, Craig SR. Methylmercury production in the anoxic hypolimnion of a dimictic seepage lake. Water, Air, and Soil Pollution. 1995;80:735–745.

9. Compeau GC, Bartha R. Sulfate-reducing bacteria: Principal methylators of mercury in anoxic estuarine sediment. Appl Environ Microbiol. 1985;50(2):498–502.

10. Gilmour CC, Henry EA, Mitchell R. Sulfate stimulation of mercury methylation in freshwater sediments. Environ Sci Technol. 1992;26(11):2281–2287. doi:10.1021/es00035a029

11. Eckley CS, Watras CJ, Hintelmann H, Morrison K, Kent AD, Regnell O. Mercury methylation in the hypolimnetic waters of lakes with and without connection to wetlands in northern Wisconsin. Can J Fish Aquat Sci. 2005;62(2):400–411. doi:10.1139/f04-205

12. Jones DS, Walker GM, Johnson NW, Mitchell CPJ, Coleman Wasik JK, Bailey JV. Molecular evidence for novel mercury methylating microorganisms in sulfate-impacted lakes. ISME J. February 2019. doi:10.1038/s41396-019-0376-1

13. Watras CJ, Bloom NS. The vertical distribution of mercury species in Wisconsin lakes: Accumulation in plankton layers. In: Mercury Pollution: Integration and Synthesis. Lewis Publications; 1994:137-151.

14. Graham AM, Aiken GR, Gilmour CC. Dissolved organic matter enhances microbial mercury methylation under sulfidic conditions. Environ Sci Technol. 2012;46(5):2715–2723. doi:10.1021/es203658f

15. Hsu-Kim H, Kucharzyk KH, Zhang T, Deshusses MA. Mechanisms regulating mercury bioavailability for methylating microorganisms in the aquatic environment: A critical review. Environ Sci Technol. 2013;47(6):2441–2456. doi:10.1021/es304370g

16. Warner KA, Roden EE, Bonzongo J-C. Microbial mercury transformation in anoxic freshwater sediments under iron-reducing and other electron-accepting conditions. Environ Sci Technol. 2003;37(10):2159–2165. doi:10.1021/es0262939

17. Hamelin S, Amyot M, Barkay T, Wang Y, Planas D. Methanogens: Principal methylators of mercury in lake periphyton. Environ Sci Technol. 2011;45(18):7693–7700. doi:10.1021/es2010072

18. Kerin EJ, Gilmour CC, Roden E, Suzuki MT, Coates JD, Mason RP. Mercury methylation by dissimilatory iron-reducing bacteria. Appl Environ Microbiol. 2006;72(12):7919–7921. doi:10.1128/AEM.01602-06

19. Guimarães JRD, Mauro JBN, Meili M, et al. Simultaneous radioassays of bacterial production and mercury methylation in the periphyton of a tropical and a temperate wetland. Journal of Environmental Management. 2006;81(2):95–100. doi:10.1016/j.jenvman.2005.09.023

20. Parks JM, Johs A, Podar M, et al. The genetic basis for bacterial mercury methylation. Science. 2013;339(6125):1332–1335. doi:10.1126/science.1230667

21. Gilmour CC, Podar M, Bullock AL, et al. Mercury Methylation by Novel Microorganisms from New Environments. Environ Sci Technol. 2013;47(20):11810–11820. doi:10.1021/es403075t

22. Gilmour CC, Bullock AL, McBurney A, Podar M, Elias DA. Robust Mercury Methylation across Diverse Methanogenic *Archaea*. mBio. 2018;9(2):1–13. doi:10.1128/mBio.02403-17

23. McDaniel EA, Peterson BD, Stevens SLR, Tran PQ, Anantharaman K, McMahon KD. Expanded phylogenetic diversity and metabolic flexibility of microbial mercury methylation. bioRxiv. [preprint](2020-01). doi:10.1101/2020.01.16.909358

24. Podar M, Gilmour CC, Brandt CC, et al. Global prevalence and distribution of genes and microorganisms involved in mercury methylation. Sci Adv. 2015;1:1–12. doi:10.1126/sciadv.1500675

25. Villar E, Cabrol L, Heimbürger-Boavida L. Widespread microbial mercury methylation genes in the global ocean. Environ Microbiol Reports. February 2020:1–11. doi:10.1111/1758-2229.12829

26. Bae H-S, Dierberg FE, Ogram A. Syntrophs dominate sequences associated with the mercury methylation-related gene *hgcA* in the Water Conservation Areas of the Florida Everglades. Appl Environ Microbiol. 2014;80(20):6517–6526. doi:10.1128/AEM.01666-14

27. Christensen GA, Wymore AM, King AJ, et al. Development and validation of broad-range qualitative and clade-specific quantitative molecular probes for assessing mercury methylation in the environment. Appl Environ Microbiol. 2016;82(19):6068–6078. doi:10.1128/AEM.01271-16

28. Gionfriddo CM, Wymore AM, Jones DS, et al. An improved *hgcAB* primer set and direct high-throughput sequencing expand Hg-methylator diversity in nature. bioRxiv. [preprint](2020-03-11). doi:10.1101/2020.03.10.983866

29. Liu Y-R, Yu R-Q, Zheng Y-M, He J-Z. Analysis of the microbial community structure by monitoring an Hg methylation gene (hgcA) in paddy soils along an Hg gradient. Appl Environ Microbiol. 2014;80(9):2874–2879. doi:10.1128/AEM.04225-13

30. Schaefer JK, Kronberg R-M, Morel FMM, Skyllberg U. Detection of a key Hg methylation gene, *hgcA*, in wetland soils: Detection of the Hg methylation gene, hgcA, in soils. Environ Microbiol Reports. 2014;6(5):441–447. doi:10.1111/1758-2229.12136

31. Bouchet S, Goñi-Urriza M, Monperrus M, et al. Linking microbial activities and low-molecular-weight thiols to Hg methylation in biofilms and periphyton from high-altitude tropical lakes in the Bolivian Altiplano. Environ Sci Technol. 2018;52(17):9758–9767. doi:10.1021/acs.est.8b01885

32. Bravo AG, Zopfi J, Buck M, et al. Geobacteraceae are important members of mercury-methylating microbial communities of sediments impacted by waste water releases. ISME J. 2018;12(3):802–812. doi:10.1038/s41396-017-0007-7

33. Christensen GA, Gionfriddo CM, King AJ, et al. Determining the Reliability of Measuring Mercury Cycling Gene Abundance with Correlations with Mercury and Methylmercury Concentrations. Environ Sci Technol. 2019;53(15):8649–8663. doi:10.1021/acs.est.8b06389

34. Liu Y-R, Johs A, Bi L, et al. Unraveling microbial communities associated with methylmercury production in paddy soils. Environ Sci Technol. 2018;52(22):13110–13118. doi:10.1021/acs.est.8b03052

35. Eddy SR. Hidden Markov Models. Current Opinion in Structural Biology. 1996;6:361–365.

36. Gionfriddo CM, Tate MT, Wick RR, et al. Microbial mercury methylation in Antarctic sea ice. Nature Microbiology. 2016;1(10). doi:10.1038/nmicrobiol.2016.127

37. Tyson GW, Chapman J, Hugenholtz P, et al. Community structure and metabolism through reconstruction of microbial genomes from the environment. Nature. 2004;428(6978):37–43. doi:10.1038/nature02340

38. Cline JD. Spectrophotometric determine of hydrogen sulfide in natural waters. Limnol Oceanogr. 1969;14(3):454–458. doi:10.4319/lo.1969.14.3.0454

39. Joshi N, Fass J. Sickle: A Sliding-Window, Adaptive, Quality-Based Trimming Tool for FastQ Files.; 2011. https://github.com/najoshi/sickle.

40. Nurk S, Meleshko D, Korobeynikov A, Pevzner PA. metaSPAdes: A new versatile metagenomic assembler. Genome Research. 2017;27(5):824–834. doi:10.1101/gr.213959.116

41. Eddy SR. Hmmer.; 2015. http://hmmer.org/.

42. Fu L, Niu B, Zhu Z, Wu S, Li W. CD-HIT: Accelerated for clustering the next-generation sequencing data. Bioinformatics. 2012;28(23):3150–3152. doi:10.1093/bioinformatics/bts565

43. Bushnell B. BBMap Short Read Aligner.; 2015. https://sourceforge.net/projects/bbmap/.

44. Hyatt D, Chen G-L, LoCascio PF, Land ML, Larimer FW, Hauser LJ. Prodigal: Prokaryotic gene recognition and translation initiation site identification. BMC Bioinformatics. 2010;11(1):119. doi:10.1186/1471-2105-11-119

45. Alneberg J, Bjarnason BS, de Bruijn I, et al. Binning metagenomic contigs by coverage and composition. Nat Methods. 2014;11(11):1144–1146. doi:10.1038/nmeth.3103

46. Kang DD, Li F, Kirton E, et al. MetaBAT 2: An adaptive binning algorithm for robust and efficient genome reconstruction from metagenome assemblies. PeerJ. 2019;7:e7359. doi:10.7717/peerj.7359

47. Sieber CMK, Probst AJ, Sharrar A, et al. Recovery of genomes from metagenomes via a dereplication, aggregation and scoring strategy. Nat Microbiol. 2018;3(7):836–843. doi:10.1038/s41564-018-0171-1

48. Wu Y-W, Simmons BA, Singer SW. MaxBin 2.0: An automated binning algorithm to recover genomes from multiple metagenomic datasets. Bioinformatics. 2016;32(4):605–607. doi:https://doi.org/10.1093/bioinformatics/btv638

49. Parks DH, Imelfort M, Skennerton CT, Hugenholtz P, Tyson GW. CheckM: Assessing the quality of microbial genomes recovered from isolates, single cells, and metagenomes. Genome Res. 2015;25(7):1043–1055. doi:10.1101/gr.186072.114

50. Eren AM, Esen ÖC, Quince C, et al. Anvi’o: An advanced analysis and visualization platform for ‘omics data. PeerJ. 2015;3:e1319. doi:10.7717/peerj.1319

51. Chaumeil P-A, Mussig AJ, Hugenholtz P, Parks DH. GTDB-Tk: A toolkit to classify genomes with the Genome Taxonomy Database. Bioinformatics. November 2019:btz848. doi:10.1093/bioinformatics/btz848

52. Konwar KM, Hanson NW, Pagé AP, Hallam SJ. MetaPathways: A modular pipeline for constructing pathway/genome databases from environmental sequence information. BMC Bioinformatics. 2013;14(202). doi:10.1186/1471-2105-14-202

53. Anantharaman K, Brown CT, Hug LA, et al. Thousands of microbial genomes shed light on interconnected biogeochemical processes in an aquifer system. Nat Commun. 2016;7(13219):1–11. doi:10.1038/ncomms13219

54. Edgar RC. MUSCLE: A multiple sequence alignment method with reduced time and space complexity. BMC Bioinformatics. 2004;5(113):1–19. doi:10.1186/1471-2105-5-113

55. Criscuolo A, Gribaldo S. BMGE (Block Mapping and Gathering with Entropy): A new software for selection of phylogenetic informative regions from multiple sequence alignments. BMC Evol Biol. 2010;10(210):1–21. doi:10.1186/1471-2148-10-210

56. Stamatakis A. RAxML version 8: A tool for phylogenetic analysis and post-analysis of large phylogenies. Bioinformatics. 2014;30(9):1312–1313. doi:10.1093/bioinformatics/btu033

57. Aberer AJ, Krompass D, Stamatakis A. Pruning rogue taxa improves phylogenetic accuracy: An efficient algorithm and webservice. Systematic Biology. 2013;62(1):162–166. doi:10.1093/sysbio/sys078

58. Schliep KP. Phangorn: Phylogenetic analysis in R. Bioinformatics. 2011;27(4):592–593. doi:10.1093/bioinformatics/btq706

59. Yu G, Smith DK, Zhu H, Guan Y, Lam TT. Ggtree: An R package for visualization and annotation of phylogenetic trees with their covariates and other associated data. Methods Ecol Evol. 2017;8:28–36. doi:10.1111/2041-210X.12628

60. Stauffer RE. Cycling of manganese and iron in Lake Mendota, Wisconsin. Environ Sci Technol. 1986;20(5):449–457. doi:10.1021/es00147a002

61. Ingvorsen K, Zeikus JG, Brock TD. Dynamics of bacterial sulfate reduction in a eutrophic lake. Appl Environ Microbiol. 1981;42(6):1029–1036. doi:10.1128/AEM.42.6.1029-1036.1981

62. Ranchou-Peyruse M, Monperrus M, Bridou R, et al. Overview of mercury methylation capacities among anaerobic bacteria including representatives of the sulphate-reducers: Implications for environmental studies. Geomicrobiology Journal. 2009;26(1):1–8. doi:10.1080/01490450802599227

63. Sczyrba A, Hofmann P, Belmann P, et al. Critical assessment of metagenome interpretation—a benchmark of metagenomics software. Nat Methods. 2017;14(11):1063–1071. doi:10.1038/nmeth.4458

64. Gilmour CC, Elias DA, Kucken AM, et al. Sulfate-reducing bacterium *Desulfovibrio Desulfuricans ND132* as a model for understanding bacterial mercury methylation. Appl Environ Microbiol. 2011;77(12):3938–3951. doi:10.1128/AEM.02993-10

65. Hernsdorf AW, Amano Y, Miyakawa K, et al. Potential for microbial H2 and metal transformations associated with novel bacteria and archaea in deep terrestrial subsurface sediments. ISME J. 2017;11(8):1915–1929. doi:10.1038/ismej.2017.39

66. Kantor RS, Huddy RJ, Iyer R, et al. Genome-resolved meta-omics ties microbial dynamics to process performance in biotechnology for thiocyanate degradation. Environ Sci Technol. 2017;51(5):2944–2953. doi:10.1021/acs.est.6b04477

67. Probst AJ, Ladd B, Jarett JK, et al. Differential depth distribution of microbial function and putative symbionts through sediment-hosted aquifers in the deep terrestrial subsurface. Nat Microbiol. 2018;3(3):328–336. doi:10.1038/s41564-017-0098-y

68. Qiu Y-L, Tourlousse DM, Matsuura N, Ohashi A, Sekiguchi Y. Draft genome sequence of *Paludibacter Jiangxiensis* NM7 ^T^, a propionate-producing fermentative bacterium. Genome Announc. 2017;5(29):e00667–17, /ga/5/29/e00667-17.atom. doi:10.1128/genomeA.00667-17

69. Spring S, Bunk B, Spröer C, et al. Characterization of the first cultured representative of Verrucomicrobia subdivision 5 indicates the proposal of a novel phylum. ISME J. 2016;10(12):2801–2816. doi:10.1038/ismej.2016.84

70. Spring S, Brinkmann N, Murrja M, Spröer C, Reitner J, Klenk H-P. High diversity of culturable prokaryotes in a lithifying hypersaline microbial mat. Geomicrobiology Journal. 2015;32(3-4):332–346. doi:10.1080/01490451.2014.913095

71. Rothery RA, Workun GJ, Weiner JH. The prokaryotic complex iron–sulfur molybdoenzyme family. Biochimica et Biophysica Act. 2008;1778:1897–1929. doi:10.1016/j.bbamem.2007.09.002

72. Jiménez Otero F, Chan CH, Bond DR. Identification of different putative outer membrane electron conduits necessary for Fe(III) citrate, Fe(III) oxide, Mn(IV) oxide, or electrode reduction by *Geobacter Sulfurreducens*. J Bacteriol. 2018;200(19):e00347–18, /jb/200/19/e00347-18.atom. doi:10.1128/JB.00347-18

73. He S, Stevens SLR, Chan L-K, Bertilsson S. Ecophysiology of freshwater Verrucomicrobia inferred from metagenome-assembled genomes. mSphere. 2017;2(5):1–17.

74. Simon J. Enzymology and bioenergetics of respiratory nitrite ammonification. FEMS Microbiol Rev. 2002;26(3):285–309. doi:10.1111/j.1574-6976.2002.tb00616.x

75. Simon J, Kern M, Hermann B, Einsle O, Butt JN. Physiological function and catalytic versatility of bacterial multihaem cytochromes c involved in nitrogen and sulfur cycling. Biochemical Society Transactions. 2011;39(6):1864–1870. doi:10.1042/BST20110713

76. Das A, Silaghi-Dumitrescu R, Ljungdahl LG, Kurtz DM. Cytochrome bd oxidase, oxidative stress, and dioxygen tolerance of the strictly anaerobic bacterium *Moorella Thermoacetica*. Journal of Bacteriology. 2005;187(6):2020–2029. doi:10.1128/JB.187.6.2020-2029.2005

77. Nobu MK, Narihiro T, Rinke C, et al. Microbial dark matter ecogenomics reveals complex synergistic networks in a methanogenic bioreactor. ISME J. 2015;9(8):1710–1722. doi:10.1038/ismej.2014.256

78. Billen G. Modelling the processes of organic matter degradation and nutrients recycling in sedimentary systems. In: Sediment Microbiology (Special Publications of the Society for General Microbiology, Book 7). 1st ed. Academic Press Inc; 1982:15-52.

79. Boschker HTS. Decomposition of organic matter in the littoral sediments of a lake. 1997.

80. Meyer-Reil L-A. Seasonal and spatial distribution of extracellular enzymatic activities and microbial incorporation of dissolved organic substrates in marine sediments. Appl Environ Microbiol. 1987;53(8):1748–1755. doi:10.1128/AEM.53.8.1748-1755.1987

81. Brock TD. A Eutrophic Lake: Lake Mendota, Wisconsin. 1st ed. Springer; 1985.

82. Bertocchi C, Navarini L, Cesaro A, Anastasio M. Polysaccharides from cyanobacteria. Carbohydrate Polymers. 1990;12:127–153.

83. Beversdorf LJ, Miller TR, McMahon KD. The role of nitrogen fixation in cyanobacterial bloom toxicity in a temperate, eutrophic lake. PLoS ONE. 2013;8(2):e56103. doi:10.1371/journal.pone.0056103

